# Tissue and cell-type specific molecular and functional signatures of 16p11.2 reciprocal genomic disorder across mouse brain and human neuronal models

**DOI:** 10.1101/2022.05.12.491670

**Authors:** Derek J.C. Tai, Parisa Razaz, Serkan Erdin, Dadi Gao, Jennifer Wang, Xander Nuttle, Celine E. de Esch, Ryan L Collins, Benjamin B. Currall, Kathryn O’Keefe, Nicholas D. Burt, Rachita Yadav, Lily Wang, Kiana Mohajeri, Tatsiana Aneichyk, Ashok Ragavendran, Alexei Stortchevoi, Elisabetta Morini, Weiyuan Ma, Diane Lucente, Alex Hastie, Raymond J. Kelleher, Roy H. Perlis, Michael E. Talkowski, James F. Gusella

**Affiliations:** Psychiatric and Neurodevelopmental Genetics Unit, Center for Genomic Medicine, Massachusetts General Hospital, Boston, MA 02114, USA; Molecular Neurogenetics Unit, Center for Genomic Medicine, Massachusetts General Hospital, Boston, MA 02114, USA; Department of Neurology, Massachusetts General Hospital and Harvard Medical School, Boston, MA 02114, USA; Program in Medical and Population Genetics, Broad Institute of MIT and Harvard, Cambridge, MA 02142, USA; Center for Quantitative Health, Division of Clinical Research and Center for Genomic Medicine, Massachusetts General Hospital, Boston, MA 02114; Department of Psychiatry, Massachusetts General Hospital and Harvard Medical School, Boston, MA 02114, USA; Bionano Genomics, San Diego, CA 92121, USA; Stanley Center for Psychiatric Research, Broad Institute of MIT and Harvard, Cambridge, MA 02142, USA; Department of Genetics, Blavatnik Institute, Harvard Medical School, Boston, MA 02115, USA; Harvard Stem Cell Institute, Harvard University, Cambridge, MA 02138, USA

**Keywords:** Genomic disorder, copy number variation, transcriptome, RNAseq, 16p11.2, CRISPR, cerebral organoid

## Abstract

Recurrent deletion and duplication of ∼743 kilobases of unique genomic sequence and segmental duplications at chromosome 16p11.2 underlie a reciprocal genomic disorder (RGD; OMIM 611913 and 614671) associated with neurodevelopmental and psychiatric phenotypes, including intellectual disability, autism spectrum disorder (ASD), and schizophrenia (SCZ). To define molecular alterations associated with the 16p11.2 RGD, we performed transcriptome analyses of mice with reciprocal copy number variants (CNVs) of the syntenic chromosome 7qF3 region and human neuronal models derived from isogenic human induced pluripotent stem cells (hiPSCs) carrying CRISPR-engineered CNVs at 16p11.2. Analysis of differentially expressed genes (DEGs) in mouse cortex, striatum, cerebellum and three non-brain tissues, as well as in human neural stem cells and induced glutamatergic neurons revealed that the strongest and most consistent effects occurred within the CNV sequence, with notable instances of differential expression of genes in the immediate vicinity that could reflect position effect. While differential expression of genes outside of chromosome 16p11.2 was largely region, tissue, and cell type-specific, a small but significant minority of such DEGs was shared between brain regions or human cell types. Gene Ontology (GO) enrichment analyses to identify cellular processes dysregulated due to these CNVs found support in select circumstances for terms related to energy metabolism, RNA metabolism, and translation but did not reveal a single universally affected process. Weighted gene co-expression network analysis identified modules that showed significant correlation with reciprocal or individual CNV genotype and better captured shared effects, indicating that energy metabolism, RNA metabolism, translation and protein targeting were disrupted across all three brain regions. The first two of these processes also emerged in the human neural stem cell (NSC) data. A subset of co-expression modules that correlated with CNV genotype revealed significant enrichments for known neurodevelopmental disorder genes, loss-of-function constrained genes, FMRP targets, and chromatin modifiers. Intriguingly, neuronal differentiation of the hiPSCs revealed that both the deletion and duplication CNV resulted in similar deficits in neurite extension and branching and alterations in electrical activity. Finally, generation of cerebral organoid derivatives indicated that the CNVs reciprocally altered the ratio of excitatory and inhibitory GABAergic neurons generated during *in vitro* neurodevelopment, consistent with a major mechanistic hypothesis for ASD. Collectively, our data suggest that the 16p11.2 RGD involves disruption of multiple biological processes, with a relative impact that is context-specific. Perturbation of individual and multiple genes within the CNV region will be required to dissect single-gene effects, uncover regulatory interactions, and define how each contributes to abnormal neurodevelopment.

## INTRODUCTION

Reciprocal genomic disorders (RGDs) are syndromes caused by recurrent CNVs generated from non-allelic homologous recombination (NAHR) (Gu et al., 2008). These disorders typically involve altered dosage of multiple genes and are collectively among the greatest contributors to neurodevelopmental disorders (NDDs) and a spectrum of related neuropsychiatric disorders (Cooper et al., 2011; Geschwind and Flint, 2015; Girirajan et al., 2011; Sebat et al., 2007; Stefansson et al., 2008). Despite this considerable morbidity, the molecular mechanisms by which these reciprocal rearrangements disrupt development remain largely unknown. Given that NAHR reproducibly alters the dosage of precisely the same sets of genes, the inherent genomic architecture of RGDs has largely prevented assessment of each affected gene’s specific contributions to associated phenotypes. However, the establishment of accessible RGD mouse models, and recent advances in CRISPR-based genome engineering in hiPSCs show promise that dissecting specific genetic and molecular underpinnings of RGDs may now be tractable (Tai et al., 2016).

Recurrent deletion and duplication of an ∼743 kb genomic segment of chromosome (chr) 16p11.2 underlies a relatively common and highly penetrant RGD associated with a spectrum of phenotypes, including autism spectrum disorder (ASD), schizophrenia (SCZ), abnormal head circumference, altered body mass, craniofacial and skeletal anomalies, and predisposition to neuroblastoma7-18. This specific CNV includes a unique ∼593 kb segment, as well as at least one copy-equivalent of an ∼150 kb flanking segmental duplication (SD). The unique segment encompasses 27 protein-coding genes while four protein-coding gene paralogues are located in each SD (Figure 1A). Genes in this region exhibit conserved-order synteny on mouse chr 7qF3, with three of the genes in the SD being present but not duplicated and the fourth being absent from the mouse genome. The 16p11.2 genes are involved in a wide variety of cellular functions, including chromatin remodeling (*INO80E, HIRIP3*) (Ding et al., 2018; Lorain et al., 1998), ubiquitination (*KCTD13*) (Escamilla et al., 2017), DNA repair (*SLX1A/B*) (Fekairi et al., 2009), MAP kinase signaling (*MAPK3, TAOK2*) (Chen and Cobb, 2001) and neurotransmitter release (*DOC2A, PRRT2*), among others (Courtney et al., 2018; Valente et al., 2016). In large-scale exome sequencing studies of ASD or NDD, no individual genes within 16p11.2 have been implicated as contributors to these disorders based on a significant excess of de novo loss-of-function mutations. Other reports have linked individual genes to these phenotypes, including a study observing an excess of missense variants in *MAPK3* among NDD cases (Coe et al., 2019), a case report where a 118 kb deletion encompassing five genes (*MVP, CDIPT, SEZ6L2, ASPHD1, KCTD13*) segregated with ASD in a three-generation pedigree (Crepel et al., 2011), and most recently publication of a nominal association of coding variants in *CORO1A* with ASD (False Discovery Rate [FDR] q < 0.05) (Fu et al., 2021), but none of these results reach stringent statistical thresholds for reproducible association comparable to well-established ASD and NDD risk loci. Multiple in vivo studies have also suggested a contribution of reciprocally modulated expression of *KCTD13* to the neuroanatomical changes associated with the 16p11.2 RGD, but these findings have also not been consistent across studies (Arbogast et al., 2019; Blaker-Lee et al., 2012; Escamilla et al., 2017; Golzio et al., 2012; Lin et al., 2015). Thus, the precise pathogenic mechanisms associated with reciprocal dosage changes of the 16p11.2 locus and the particular genes that drive them remain to be defined.

**Figure. 1.**
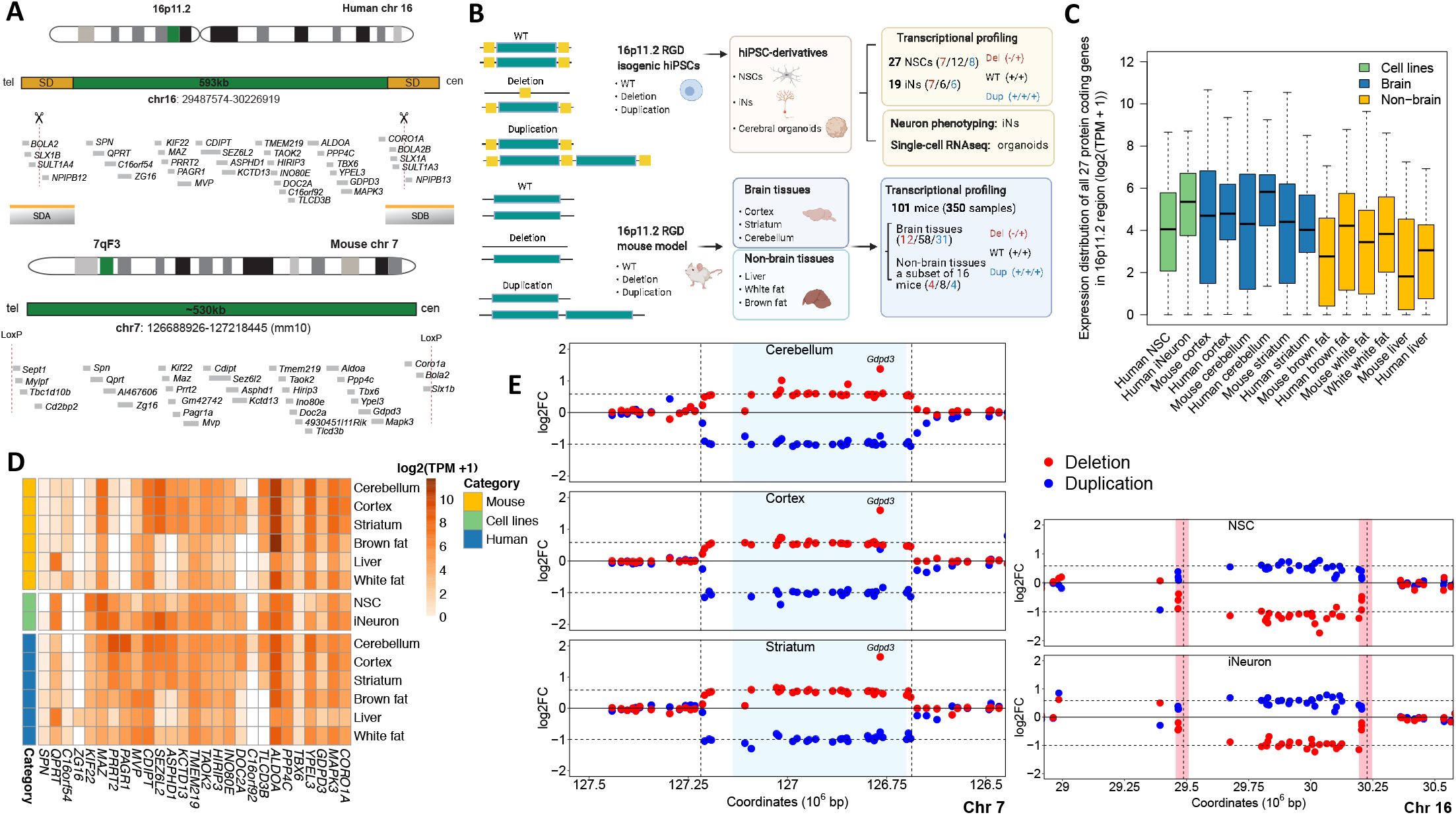
Figure 1. Experimental design and expression profile of RGD genes across samples. (A) Illustration of human 16p11.2 segment and SD and the syntenic 7qF3 region in mouse. Only protein coding genes are shown using human Ensembl GRCh37 (version 75) annotation and mouse Ensembl GRCm38 (version 83). Single-guide RNA targeting the SDs to promote a model of NAHR-mediated CNV as indicated by scissor. Mouse models were generated through Cre-loxP-mediated recombination as described (Horev et al., 2011). (B) Schematic of the study design and analyses. To systemically dissect molecular functions associated with 16p11.2 RGD, we performed transcriptome analyses of 101 mice with reciprocal CNVs (350 total samples) of the syntenic chr 7qF3 region across cortex, striatum, and cerebellum, as well as three non-brain tissues. Furthermore, we generated NSCs, iNs, and cerebral organoid derivatives of isogenic hiPSC harboring CRISPR-engineered reciprocal 16p11.2 CNVs and assessed cellular, transcriptional, and single-cell signatures associated with 16p11.2 CNV. (C) Expression distribution of 27 protein coding genes within 16p11.2 region in WT samples including mouse tissues, NSC and iNs, and GTEx data. The results revealed 16p11.2 genes with higher expression level in brain tissue and neuron than non-brain tissues. (D) Heatmaps of 16p11.2 region genes’ basal expression in mouse tissues (Top), hiPSC derived NSCs and induced neurons (Middle), and human (GTEx, transcripts-per-million) (Bottom). (E) Fold change (log2) of the protein coding genes in the CNV and in the flanking regions are shown in coordinate space for deletions in red and duplications in blue across brain tissues (left panel) and human cells (right panel). Light blue shaded region in the left panel highlights the unique portion of 16p11.2 CNV region harboring 27 human orthologous protein coding genes in the mouse 7qF3 segment, whereas pink vertical bars in the right panel highlight the segmental duplication region in the human 16p11.2 segment.

Recent genomic studies of the broad set of genes individually associated with ASD and NDD have suggested a convergence of NDD genes on key functional pathways including chromatin modification, transcriptional regulation, and synaptic transmission (De Rubeis et al., 2014; Satterstrom et al., 2020). Investigations of animal models and tissues have also explored NDD pathogenesis by interrogating protein-protein interactions, quantifying regional and temporal patterns of co-expression (Grove et al., 2019; Krishnan et al., 2016; Parikshak et al., 2013; De Rubeis et al., 2014; Satterstrom et al., 2020; Willsey et al., 2013), and performing in vivo phenotyping during development (Willsey et al., 2021). The resulting data have corroborated the aforementioned processes as contributors but have also revealed effects on neurogenesis that include altered ratios of cell types in the developing brain. One approach to elucidate pathogenic mechanisms in RGDs is to use global transcriptome analysis of peripheral cells from individuals harboring CNVs (Blumenthal et al., 2014). However, patient-specific variability in genetic background and cell type-specific expression patterns can complicate identification of regulatory changes most relevant to abnormal neurodevelopment. We previously developed genome editing methods to model NAHR by targeting SDs with CRISPR/ Cas9 to produce precise isogenic human cellular models of recurrent RGDs in hiPSCs and derived neuronal lineages (Tai et al., 2016). Here, we sought to integrate large-scale human and mouse modeling to disentangle the tissue-specific, cell-type specific, and gene dosage-specific molecular and transcriptional signatures associated with 16p11.2 RGD. To accomplish this, we examined tissue-specific changes in global gene expression using mouse models with reciprocal CNVs in 7qF3 (Horev et al., 2011) and comprehensive analyses of three brain regions (cortex, striatum, cerebellum) and three non-brain tissues (liver, white fat, brown fat), along with NSCs and neurogenin-2 induced neurons (iNs) derived from isogenic hiPSC lines engineered to model reciprocal 16p11.2 CNV (Figure 1B). We find that these CNVs cause a complex spectrum of distinct and overlapping gene expression changes that reflect both tissue-specific and shared pathway changes. Human cell models carrying 16p11.2 CNVs demonstrate aberrant neuronal phenotypes, including shorter neurites and reduced electrical activity. Finally, single-cell RNA sequencing (scRNAseq) of cerebral organoids harboring 16p11.2 CNVs revealed an altered cell composition with an excitatory/ inhibitory neuron imbalance, providing a potential link between 16p11.2 rearrangements and their associated neuropsychiatric phenotypes, including ASD and SCZ.

## RESULTS

### The genomic properties of the 16p11.2 RGD locus

As described above, the human 16p11.2 RGD locus involves ∼743 kb of genomic sequence that includes a unique ∼593 kb segment, as well as at least one copy-equivalent of an ∼150 kb flanking SD. The unique segment encompasses 27 protein coding genes while four gene paralogues are located in each SD (Figure 1A). Across these 31 protein coding genes in humans, 30 (with the exception of *ZG16*) are expressed at detectable levels (≥ 0.1 TPM) from bulk RNA-sequencing (RNAseq) across at least one of 13 brain regions profiled in the Genotype-Tissue Expression (GTEx) project, with the highest fraction of genes expressed in the frontal cortex (30/31, 97%) (Lonsdale et al., 2013). To place the genomic properties of the 16p11.2 RGD, and the genes included within, in the context of 19 NAHR-mediated RGDs profiled in a study by Collins et al. (2021) in which both deletion and duplication were associated with a spectrum of disease phenotypes, the proximal 16p11.2 CNV ranked fifth for total number of protein-coding genes among all RGD segments but ranked first after normalizing by RGD size (i.e., it is the most gene dense RGD profiled). The 16p11.2 RGD also ranked fourth for normalized density of constrained genes intolerant to loss-of-function (LoF) variation as defined as the bottom (decile or sextile) of the LoF observed over expected upper bound fraction (LOEUF) metric in the Genome Aggregation Database (gnomAD) (Karczewski et al., 2020). When we further scrutinized these 19 NAHR mediated GD segments for more nuanced metrics of dosage sensitivity (i.e., intolerance to haploinsufficiency [pHaplo] or triplosensitivity [pTriplo]) provided in Collins et al. (2021), the 16p11.2 CNV ranked second in terms of the combined number of predicted haploinsufficient and/or triplosensitive genes when normalized by GD size. These data collectively suggest that the functional consequences of reciprocal 16p11.2 CNV are likely due to a number of dosage sensitive loci and not concentrated for all phenotypes on a single ‘driver gene’ like those in regions such as 15q11-13 (*UBE3A* in Angelman syndrome (Kishino et al., 1997)) or 17q21 (*KANSL1* in Koolen-de Vries syndrome (Koolen et al., 2012)), among many others.

### Transcriptional profiling of 16p11.2 RGD in mouse and human models

To identify the transcriptional consequences of the 16p11.2 CNVs in the brain, we used both mouse models of deletion and duplication of the 7qF3 region of synteny conservation and neuronal derivatives of 16p11.2 RGD hiPSC models (Figure 1 A and B). These mice display 16p11.2-relevant phenotypes due to CNVs that include orthologues of the 27 unique 16p11.2 protein-coding genes and of two genes in the flanking human segmental duplication (*SLX1A, BOLA2*), which are not duplicated in the mouse (Horev et al., 2011). The reciprocal mouse CNVs also include four genes located outside the human segment: *Cd2bp2, Tbc1d10b, Mylpf* and *Sept1*. We performed transcriptional profiling in brain on a large collection of 101 mice (12 deletion, 31 duplication, 58 wild-type (WT) littermates) across cortex, striatum, and cerebellum (302 total libraries from brain tissue) and examined three non-brain tissues of relevance to 16p11.2 RGD phenotypes (liver, white fat, brown fat) in a subset of 16 mice (4 deletion, 4 duplication, 8 WT littermates; 48 total libraries from non-brain tissue). In sum we profiled tissue-specific expression patterns from mouse 7qF3 CNV across 350 RNAseq libraries (Table S1).

For the comparison with human cellular models, we generated isogenic hiPSC lines with 16p11.2 CNVs using the CRISPR SCORE method and genotyped them by quantitative real-time PCR (qRT-PCR) and genome-wide array-based comparative genomic hybridization (aCGH). For a subset of lines, we additionally performed nanopore sequencing and generated Direct Label and Stain (DLS) optical genome maps (Bionano Genomics, CA) (Figure S1A, see Methods). We then differentiated the hiPSCs into NSCs (n= 27 lines; 12 WT, 7 deletion, 8 duplication), iNs (n=19 lines; 6 WT, 7 deletion, 6 duplication), and cerebral organoids (n=8) to assess disease-relevant cellular and transcriptomic signatures (Figure 1B). Cellular identities of all lines were verified using cell type-specific marker gene expression in the RNAseq data (Figure S1B).

We first reviewed the local expression patterns of genes within and near the RGD segment by CNV genotype. The expression levels of genes within the CNV interval in WT mice varied widely by tissue. In general, their expression levels and the significance of altered expression caused by the CNV were relatively greater in brain tissues, with some exceptions such as *Qprt*, which was predominantly expressed in the liver (Figure 1 C and D, Table S2). GTEx data show a comparable overall pattern where 16p11.2 CNV genes are more highly expressed in human brain tissues than in non-brain tissues and the human NSCs and iNs showed brain-like expression, except for *QPRT* (Figure 1 C and D). In both mouse and human experiments, expression changes of most genes within the CNV segment reflected their dosage loss or gain, as expected (Figures 1E and S1C). We found no consistent evidence in the brain regions of dosage compensation from the unaltered allele, as we had observed previously for human lymphoblastoid cells (Blumenthal et al., 2014). The SD genes *NPIPB12/NPIPB13* are members of a dispersed set of paralogues (absent in the mouse genome) that precludes their individual quantification while the paralogue pairs *BOLA2/BOLA2B, SLX1A/ SLX1B* and *SULT1A3/SULT1A4* each constitute four copy-equivalents in the WT human lines. The human deletion and duplication cell lines lost and gained one SD copy, respectively, corresponding to expected expression fold changes (FCs) of 0.75 and 1.25 for these three paralogue pairs. The average expression fold change of these SD genes in human cell lines did not deviate significantly from these expectations (iN deletion, FC = 0.79 ± 0.07, p = 0.17; iN duplication, FC = 1.25 ± 0.04, p = 0.84; NSC duplication, FC = 1.18 ± 0.11, p = 0.18), except that SD genes were significantly more down regulated than expected in the NSC deletion samples (FC = 0.64 ± 0.10, p = 0.047). A few genes in mouse tissues and human cell lines did not show altered expression levels consistent with an expected CNV effect. *Gdpd3* exhibited highly variable expression across all six mouse tissues, previously reported behavior attributed to genetic background differences of the parental mouse strains (Horev et al., 2011). Genes with higher-than-expected dysregulation include *Kif22* in brown fat, *Mylpf* in liver, *Qprt* and *Zg16* in white fat (Figure S1C), and *TLCD3B* in human NSCs (6.6-fold, Figure 1E). Overall, genes in the engineered CNV region were expressed more variably in the non-brain tissues than in the brain tissues. Principal components analysis based on the expression profile of the 27 CNV genes separated all tissues (Figure S1D).

### Genome-wide transcriptional changes in the RGD models

We next asked whether genes outside the 16p11.2 region showed altered expression patterns in our RGD models. In the mouse models, the number of differentially expressed genes (DEGs; based on FDR <0.1) due to 7qF3 deletion or duplication (excluding the CNV genes) varied greatly across tissues (Figure 2A, Table S3). While most of these effects were tissue-specific, there was significantly greater sharing of DEGs than expected by chance across the brain tissues (Figure 2A), sharing that was evident for both for deletion- and duplication-associated DEGs (Figures S2A and S2B). Conversely, within each brain region there were fewer DEGs shared between deletion and duplication models (Figure S2C). Interestingly, many of these DEGs were similarly upregulated or downregulated in both CNV models (i.e., perturbed non-reciprocally). It is possible that for some genes near the CNV region (e.g., *Ccdc101, Kdm8*) differential expression could be due to position effects observed in brain tissues but not in peripheral tissues. Only one gene, *Kctd21*, was differentially expressed due to both 7qF3 CNVs in all three brain tissues, with deletion being associated with upregulation and duplication with downregulation (i.e., reciprocal dysregulation). The overall pattern of gene expression changes observed in brain was not observed in peripheral tissues, where there was little sharing of DEGs across liver, brown fat and white fat or between these tissues and the brain regions (Figure 2A). With the notable exception of liver, these peripheral tissues, like the brain tissues, lacked significant DEG sharing between deletion and duplication models, (Figure S2C). Overall, our mouse DEG analyses highlight significant shared effects of the CNVs that are greatest in the brain but also reveal many more distinct tissue-specific reciprocal and non-reciprocal impacts.

**Figure 2.**
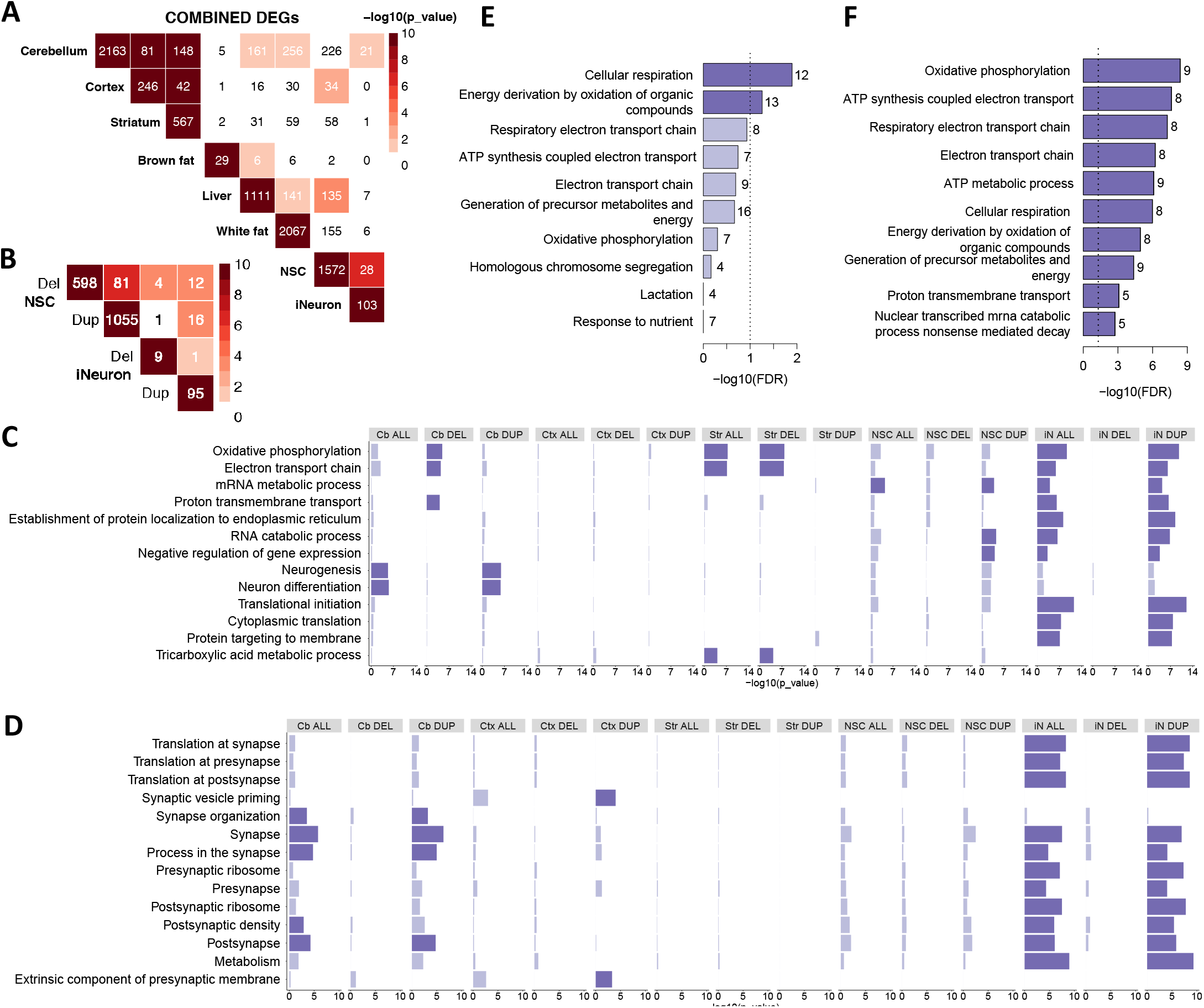
Transcriptional profiling and disease specific brain signatures in 16p11.2 transgenic mouse and human neuron models. (A) Overlap among the DEGs observed between mouse brain tissues, peripheral tissues, and human cells. (B) Overlap among the DEGs observed between NSC and iN. (C) GO enrichment analysis is shown as bar plots for mouse brain tissues, human NSCs and iNs comparisons. Enrichments at nominal level and at FDR < 0.1 are marked in light blue and blue, respectively. (D) SynGO enrichment analysis for mouse brain tissues, human NSCs and iNs comparisons. Enrichments at nominal level (p < 0.05) and at FDR < 0.1 are marked in light blue and blue, respectively. (E) The pathways enriched for shared mouse brain DEGs. (F) GO Biological Process terms enriched for DEGs shared between human NSCs and iNs.

In the human cell models, the number of global gene expression changes varied strikingly between NSCs and iNs (Figure 2A and 2B, Table S3), with the former yielding many more significant DEGs due to either 16p11.2 deletion or duplication. Despite the surprising paucity of iN DEGs, there was significant sharing of DEGs between NSC and iNs (p = 7.55e-7), although these effects were less significant than the stronger sharing exhibited across the mouse brain regions (p < 1e-13) (Figures 2A, S2A and S2B). However, the NSC DEGs showed more significant evidence than the brain regions of sharing between deletion and duplication effects (p = 6.12e-7), although only 31 of the 81 shared DEGs exhibited reciprocally altered expression (Figure 2B, Figure S2C). Thus, like the adult tissues of the mouse model, the human models of NSC and maturing neuron developmental stages point to a combination of shared and nonshared effects across cell types, with evidence for both reciprocal and non-reciprocal changes in gene expression.

### Gene Ontology enrichment analysis of differentially expressed genes

To explore the potential functional ramifications of differential expression in these models, we performed Gene Ontology (GO) Biological Process term enrichment (Figure 2C and Table S4). No GO terms were enriched at an FDR-significant level (FDR < 0.1) among mouse cortex DEGs, but the mouse cerebellar and striatal deletion-elicited DEGs both showed significant enrichments for a number of GO terms related to energy metabolism, such as oxidative phosphorylation and electron transport chain. Although the significant terms due to deletion largely overlapped between these two brain regions, a few were tissue-specific, including proton transmembrane transport in cerebellum and tricarboxylic acid cycle in striatum. In cerebellum, the duplication CNV elicited a much larger number of DEGs, which revealed enrichments for a number of terms related to neuronal development, including neuron differentiation and neurogenesis (Figure 2C). Therefore, we repeated our enrichment analyses using expert-curated SynGO database which was developed for studying synaptic biology and includes a number of additional GO Biological Process and Cellular Components terms (Figure 2D and Table S5). Notably, known NDD genes are enriched for SynGO terms, suggesting that leveraging this database may provide insights into pathogenic mechanisms. The cerebellum 7qF3 duplication DEGs showed significant enrichments for a series of terms related to the synapse, including synapse, synapse organization, process in the synapse, and postsynapse. In contrast, cortex 7qF3 duplication DEGs were enriched for synaptic vesicle priming and extrinsic component of presynaptic membrane (Figure 2D).

Given the significant sharing of DEGs across brain regions, we also performed GO term enrichment analysis for the 223 unique DEGs shared by at least two brain regions. These showed significant enrichment only for the energy metabolism-related terms cellular respiration and energy derivation by oxidation of organic compounds (Figure 2E, Table S4), both at lower significance levels than the distinct energy metabolism-related terms enriched among cerebellum and striatum DEGs (Figure 2C, Table S4). These data indicate that many additional DEGs that are not shared across the brain regions contribute to the increased significance of the energy metabolism-related terms in individual brain regions, suggesting that analysis focused solely on shared effects across tissues does not fully capture the extent of a biological process altered due to the CNV.

Consistent with the findings from the mouse brain regions, the iN DEGs associated with duplication showed FDR-significant GO term enrichment for energy metabolism-related terms, among many others (Figure 2C, Table S4). The iN duplications also revealed significance for a variety of terms related to translation, mRNA metabolism and protein targeting and some of the RNA metabolism terms (RNA catabolic process, RNA metabolic process) were shared with NSC duplication DEGs, although the latter showed fewer GO enrichments overall. Not surprisingly, the SynGO enrichment analyses of DEGs from the iNs and NSCs differed substantially, with significant terms resulting only with the more differentiated iNs, where SynGO:metabolism was prominent, along with translation at both presynapse and postsynapse (Figure 2D, Table S5). Several of the significant terms overlapped with the mouse brain analysis of cerebellar duplication-elicited DEGs. Interestingly, the 81 genes shared between 16p11.2 deletion and duplication NSCs did not detect any significant GO terms, failing to support a common process disrupted by the reciprocal dosage changes in this cell type. By contrast, analysis of the 28 DEGs shared between the NSC and iN DEGs yielded significance for a series of energy metabolism-related terms (Figure 2F, Table S4) and for terms related to synaptic protein translation (Figure S2D, Table S5) pointing to these processes as disrupted across both cell types.

None of the GO terms from brain tissue analyses emerged as significant in the peripheral tissues (Figure S3A and S3B). Brown fat DEGs revealed no significant GO term enrichment while the deletion- and duplication-elicited DEGs each yielded a distinct set of significant enrichments in liver and white fat. Liver deletion DEGs showed top enrichments for a series of terms related to muscle/actin-based contraction and filament sliding, monocarboxylic acid metabolism, and lipid metabolism, while those due to duplication revealed terms such as connective tissue development, chondrocyte differentiation, and apoptotic process. In white fat, the deletion DEGs detected most prominently a series of terms related to organic acid metabolism while the duplication DEGs revealed terms related to inflammation.

Overall, the mouse DEG and enrichment analyses suggest that in the brain, the CNVs produce both shared effects, whose strength varies greatly across the three brain regions, and effects that are largely region-specific. In the peripheral tissues, the significant differences are largely tissue- and dosage-specific. Analyses of the human 16p11.2 models reinforce the view that, at the level of FDR-significant differences, the pattern of gene expression due to the reciprocal CNVs is largely cell-type specific even when it involves the same biological process, and that disruption of genes involved in energetics and synapse-related functions is

### Co-expression analyses of mouse tissues and human cell lines

The finding that similar biological processes are revealed in different tissues by largely distinct sets of CNV-elicited DEGs indicates that an analysis limited to FDR-significant DEGs does not adequately capture the biological impact of the RGD. Consequently, as a complementary route that utilized all of the gene expression data in defining the processes disrupted by the CNVs, we performed weighted gene correlation network analysis (WGCNA) (Langfelder and Horvath, 2008a). In view of the largely tissue- and region-specific pattern of DEGs and the forced differential expression of the genes contained in the CNV region, we applied WGCNA individually to each brain region, peripheral tissue, and human cell type after excluding the 16p11.2 CNV genes. For a given analysis, we combined deletion, duplication, and wild-type datasets, performed WGCNA, and then examined the eigengenes of the resulting modules for fit to 1) a reciprocal CNV effect (CNV dosage), 2) an effect driven solely by deletion (Del vs. Dup+WT) or by duplication (Dup vs. Del+WT), or 3) a similar effect induced by both deletion and duplication (Genotype vs. WT). Within each co-expression module that showed a significant fit at p < 1e-5, we tested the member genes for enrichment of GO Biological Process terms and SynGO terms as well as enrichment of a variety of neurodevelopment-associated gene sets. The results for mouse brain regions and human cells are shown in Figure 3, and for mouse peripheral tissues are shown in Figure S3C. The latter yielded only one significant co-expression module in the liver and none in brown fat and white fat. The full lists of co-expression modules in the mouse brain and human cell data are shown in Tables S6a and S6b, respectively. Module eigengenes of all the modules identified by WGCNA for six mouse tissues and human NSCs and iNs are shown in Table S7.

**Figure 3.**
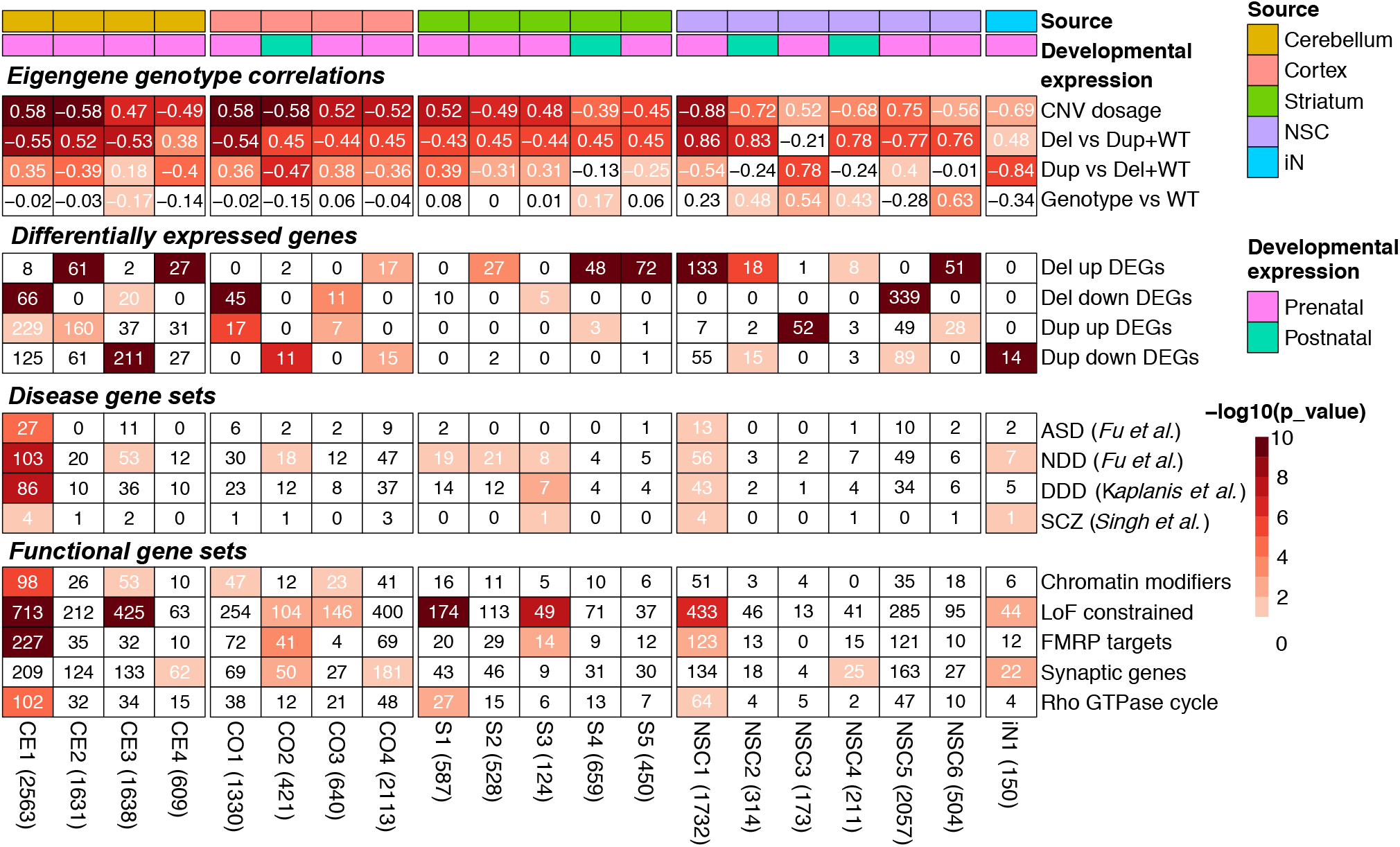
Weighted gene co-expression network analysis (WGCNA) of mouse brain and human cell transcriptomes. Modules that are statistically significantly associated with deletion (Del) and duplication (Dup) genotypes at p < 1e-5. The first and second row annotations indicate for each column the source tissue or cell line for modules, and whether module genes are more highly expressed in prenatal or postnatal stages. The first panel, named “Eigengene genotype correlations”, is colored to show the statistical significance of the listed module eigengene correlation. The second panel named “Differentially expressed genes” is colored to show the statistical significance of the overlap between module members and up- and down-differentially expressed genes (FDR < 0.1) from the corresponding mouse tissue or human cell line with deletion (Del) or duplication (Dup) genotype. The third and fourth panels list the number of genes overlapping between the module and literature-curated gene lists (disease gene sets and functional gene sets, respectively) and are colored to show the statistical significance of the overlap. The lists include ASD associated genes (Fu et al., 2021), NDD associated genes (Fu et al., 2021), DDD associated genes (Kaplanis et al., 2020), rare variants in genes associated with SCZ (Singh et al., 2022), chromatin modifiers (Iossifov et al., 2014), loss-of-function intolerance constrained genes (LOEUF < 0.35) as reported by the genome aggregation database consortium (Karczewski et al., 2020), FMRP targets (Darnell et al., 2011), synaptic genes from SynGO v1.1 (Koopmans et al., 2019), and Rho GTPase cycle genes (Gillespie et al., 2022; Subramanian et al., 2005). The heatmap color scale of highlights the statistical significance in -log10 scale, while numbers in the heatmap cells are Pearson’s correlation coefficients (the first panel) or number of genes shared between gene sets and modules (other panels). Numbers within the parentheses next to module names show the number of protein-coding genes in the co-expression modules identified in the mouse tissue, human NSCs and iNs modules.

Analysis of the cerebellum (CE modules) and cortex (CO modules) yielded four co-expression modules each (CE1-4 and CO1-4, respectively) that showed a significant fit to one or more of the models tested, while the striatum (S modules) revealed five significant modules (S1-S5) (Figure 3). For most of these co-expression modules, the eigengenes showed the greatest significance for a continuous reciprocal effect of CNV dosage, albeit with different relative contributions from deletion and duplication, as evidenced by the lesser significance achieved for solely a deletion effect (Del vs. Dup+WT) and solely a duplication effect (Dup vs. Del+WT). Notable exceptions, such as the cerebellar CE4 module and the striatal S5 module, provided greater significance for an effect limited to deletion (only enriched for Del up-regulated DEGs). In contrast, the cortex CO2 module revealed significance related to duplication (only enriched for Dup down-regulated DEGs). Notably, no module was significant for gene expression effects driven in the same direction by both deletion and duplication. Cerebellum CE1 module whose eigengene showed a positive correlation for 16p duplication, displayed the most significant enrichment for ASD (Fu et al., 2021), NDD (Fu et al., 2021), genes identified in the Deciphering Developmental Disorder Study (DDD) (Kaplanis et al., 2020), schizophrenia (SCZ) genes (Singh et al., 2022), chromatin modifiers (Iossifov et al., 2014), loss-of-function (LoF) constrained genes (Karczewski et al., 2020), mRNA targets of fragile X retardation protein (FMRP targets) (Darnell et al., 2011), and Rho GTPase cycle genes (Gillespie et al., 2022; Subramanian et al., 2005), which was the top significantly enriched term from the REACTOME database (Gillespie et al., 2022). These disease and functional gene sets have been described previously in relation to 16p11.2 CNV genes and associated neurodevelopmental and psychiatric disorders (Cooper et al., 2011; Geschwind and Flint, 2015; Girirajan et al., 2011; Sebat et al., 2007; Stefansson et al., 2008). Other modules, including CE3, CE4, CO1-4, and S1-3, displayed selective enrichments for disease and functional gene sets. The results of co-expression enrichment indicate significant disturbance of the transcriptome caused by 16p11.2 CNV, but with effects that are variable across brain regions. The distinct impact on various brain regions may indicate a potential link between tissue-specific contributions to a spectrum of phenotypes associated with 16p11.2 RGD.

To gain insights into disease development and progression, we then compared expression patterns of member genes from co-expression modules across human brain developmental time points using the expression data from the PsychENCODE project (Li et al., 2018) (Figure S3D). Except for CO2, S4, NSC2, and NSC4, most of the modules exhibited high expression in the prenatal stage, suggesting that the impact of 16p11.2 CNV is significant in early developmental stages (Figure 3 and Figure S3D). Module CO2, highly expressed in the postnatal stage, displayed significant enrichment for LoF constrained genes (Karczewski et al., 2020), FMRP targets (Darnell et al., 2011), and synaptic genes (Koopmans et al., 2019), strongly suggesting that this module contributes mainly to the abnormalities in the cortex (Figure 3). GO term enrichment revealed biological processes associated with the various co-expression modules (Figure 3, Table S8a, Table S9). Although some mouse brain modules (CO1, CO2, S3, S5) provided comparatively weak or no support for GO enrichment (FDR > 1e-5), most pointed to multiple terms with moderate to high support (FDR < 1e-5) that were most often shared with several other modules across all three brain regions, indicating disruption some of the same processes across the brain. For example, the top-scoring term in cortex (CO4, translational termination) was also significant in CE4 and S4. The top-scoring terms in cerebellum modules CE2 and CE4 (cellular respiration and cotranslational protein targeting to membrane) were also significance in S4 and CO4, respectively. The top-scoring term in striatum (S4, mitochondrion organization) was also significant in CO4 and CE2. Most of the shared terms across the modules were related to energy metabolism (e.g., cellular respiration; oxidative phosphorylation; mitochondrion organization; aerobic respiration; ATP synthesis-coupled electron transport), to RNA metabolism (e.g., RNA processing; mRNA processing; RNA splicing; mRNA metabolic process; RNA catabolic process), to translation (e.g., peptide metabolic process; translational initiation; translational elongation; translational termination; peptide biosynthetic process) or to protein targeting (e.g., establishment of protein localization to endoplasmic reticulum; protein targeting). Other GO enrichments that were detected at FDR < 1e-05 in only a single module typically also received weaker support (1e-5 < FDR < 0.1) in some other modules, but some enrichments implicated distinctly region-specific effects (e.g., cerebellum CE1, homophilic cell adhesion via plasma membrane adhesion molecules; striatal S4, proteasomal ubiquitin-dependent protein catabolic process). Interestingly, while pointing to cell adhesion, the CE1 module, did not detect any of the energy metabolism, RNA metabolism, translation or protein targeting terms, yet it was notably enriched for disease and functional gene sets. These results suggest that the co-expression of these genes in the largest module observed (2563 genes) (Table S8a) might reflect convergence of components of multiple biological processes, thereby representing disease pathways that are not well captured by individual GO terms defined from normal biological processes.

A similar analysis of the less abundant human cell line data yielded six significant modules in NSCs (NSC1-6) and one in iNs (iN1) (Figure 3, Table S8b, Table S9), generally of smaller size and lower significance than the modules detected in mouse brain. In individual instances, these modules favored reciprocal effects (NSC1) or effects primarily due to 16p11.2 deletion (NSC2,4-6) or to duplication (NSC3, iN1). NSC1 and iN1 modules displayed common enrichments for disease and functional gene sets including NDD, SCZ, and LoF constrained genes, while iN1 yielded a distinct enrichment for synaptic genes. In GO enrichment analysis, only NSC6 yielded a significant functional category at FDR < 1e-5, and most of these were among the RNA metabolism terms noted above for the mouse model. Weaker support (1e-5 < FDR < 0.1) was obtained, among others, for a variety of terms related to cellular morphogenesis (iN1, NSC4), cell substrate adhesion (NSC1), cell-cell adhesion and neurogenesis (NSC4) and energy metabolism (NSC6).

The co-expression analysis yielded both distinct and common enrichments across tissues and cells, suggesting that there are critical genes shared by particular modules. We performed a pairwise comparison of the modules that exhibited enrichment for ASD or NDD gene sets at p < 0.1 to define shared genes and signatures (Figure S3E). Amongst these comparisons, CE1 and NSC1, both containing genes highly expressed in the prenatal stage, displayed the most significant overlap (p < 1e-10) between mouse and human cell line modules: 423 shared genes were significantly enriched for disease gene sets, chromatin modifiers (Iossifov et al., 2014), LoF constrained genes (Karczewski et al., 2020), FMRP targets (Darnell et al., 2011) and Rho GTPase cycle (Gillespie et al., 2022; Subramanian et al., 2005). Interestingly, these two modules were correlated in opposite directions with CNV dosage, suggesting that some genes that impact the system during the early neurodevelopmental stages can respond to perturbation by deletion and duplication in a context-specific manner (Figure S3E).

SynGO analysis of the mouse brain and human cell WGCNA modules revealed fewersignificantterms overall at FDR< 0.1 (Table S9). In the mouse brain, these were limited to CE2 and CE4, with the latter revealing the greatest significance. However, again CE1 differed, with its top terms being related to synaptic organization and function (maintenance of alignment of postsynaptic density and presynaptic active zone; postsynaptic spectrin-associated cytoskeleton organization; regulation of presynaptic membrane potential) while the most significant enrichments in both CE2 and CE4 related to translation (e.g., SynGO: postsyn_ribosome postsynaptic ribosome, SynGO:presyn_ribosome presynaptic ribosome, GO: translation at synapse). The NSC and iNs shared enrichment of terms related to synaptic organization (e.g., synapse organization), while the top hits in each were anchored component of presynaptic active zone membrane and SynGO:synprocess process in the synapse, respectively.

### The neurons with 16p11.2 RGD display aberrant spatiotemporal neurite dynamics

To determine whether the changes in gene expression caused by the 16p11.2 deletion and duplication lesions were reflected in the functional properties of neurons carrying these lesions we evaluated whether the 16p11.2 RGD alters the morphological and electrophysiological properties of iNs. To assess neurite dynamics, we performed morphological analysis of the 16p11.2 RGD iNs using the IncuCyte real-time live-cell imaging system (Sartorius) over seven days (Figure 4A) in comparison with WT and cells heterozygous for inactivation of *KCTD13*, a candidate driver gene in the 16p CNV. Figure 4B shows the corresponding neurite outgrowth images with image segmentation. One-way ANOVA indicated significant differences in cumulative neurite length (F=25.14, p=4.12e-15, df=3) and neurite branchpoints (F=33.14, p=1.81e-19, df=3) across WT, 16p11.2 deletion (16pDel), 16p11.2 duplication (16pDup), and *KCTD13* heterozygous deletion (KCTD13Het) lines. The iNs with 16p11.2 CNVs showed lower neurite length and reduced numbers of neurite branchpoints (Figure 4CD) while the WT and KCTD13Het iNs displayed comparable values (Figure 4CD). Post hoc comparisons with the Tukey HSD test of the 16pDel (neurite length: mean=140.30, SE=7.52; neurite branchpoints: mean=3.10, SE=0.25) and 16pDup (neurite length: mean=139.68, SE=7.98; neurite branchpoints: mean=3.79, SE=0.26) iNs confirmed significantly decreased total neurite length (p=4.31e-9 and p=5.37e-9, respectively, Figure 4E) and branchpoints (p=3.77e-9 and p=4.85e-9, respectively, Figure 4F) when compared to the WT (neurite length: mean=203.73, SE=6.27; neurite branchpoints: mean=5.77, SE=0.21). The 16p11.2 CNV iNs were also significantly different from the KCTD13Het iNs (neurite length: mean=207.24, SE=8.76; neurite branchpoints: mean=6.01, SE=0.29) in terms of both neurite length (p=4.44e-8 and p=7.47e-8, respectively, Figure 4E) and neurite branchpoints (p=3.77e-9 and p=1.14e-7, respectively, Figure 4F). The 16p11.2 CNV iNs were not significantly different from each other, and there was no significant difference between the WT and KCTD13Het iNs. Taken together, these findings strongly suggest that 16p11.2 CNV results in neurite outgrowth and branching deficits, that some of the genes in the region are involved in these mechanisms, and that deletion of *KCTD13* alone does not recapitulate these neuronal deficits.

**Figure 4.**
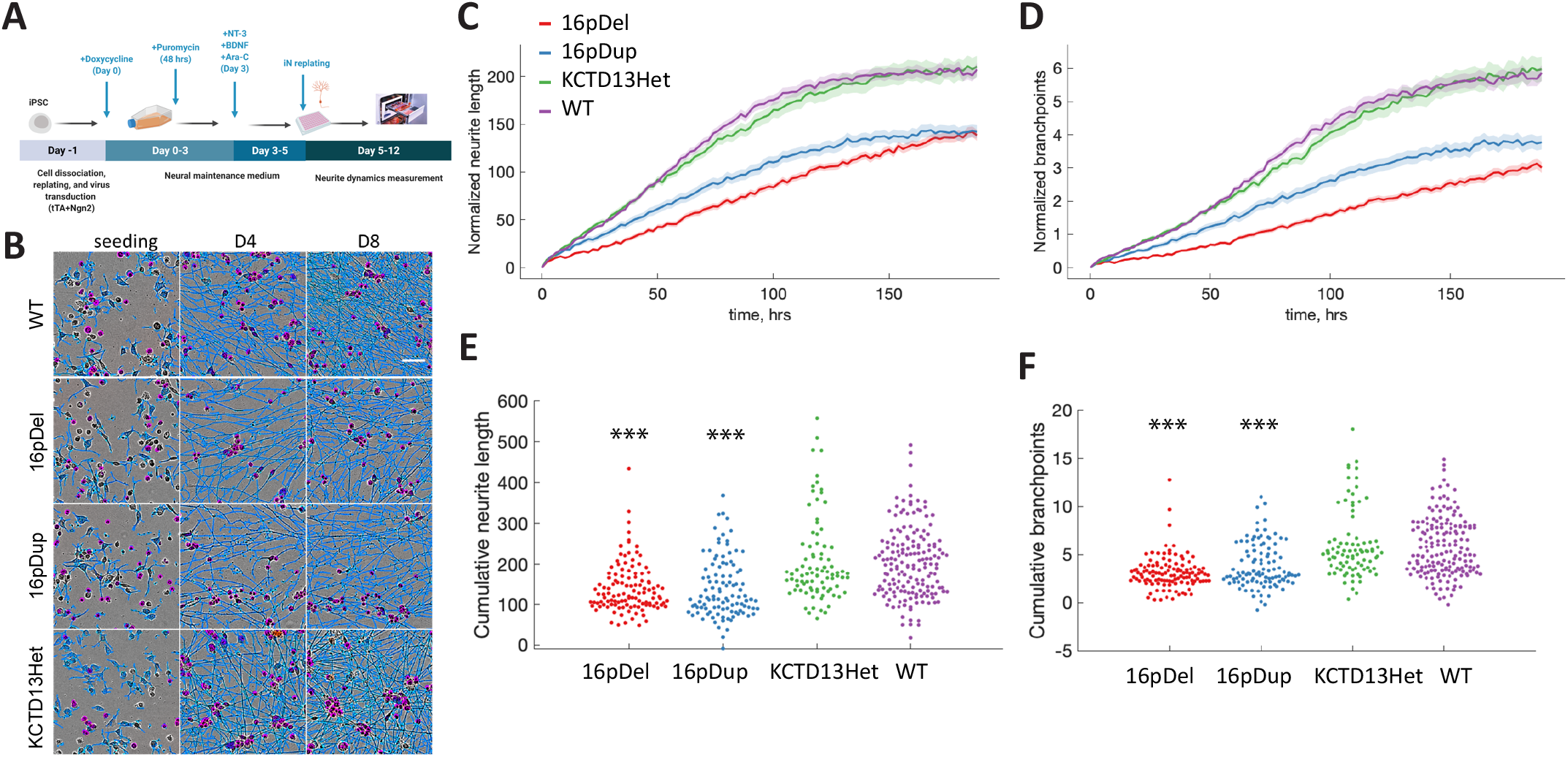
16p11.2 RGD neurons revealed altered neurite dynamic features. (A) Experimental design. iNs were differentiated from hiPSCs as described in Materials and Methods. On Day 5 iNs were replated onto 96-well plates and imaged over 7 days using Incucyte ZOOM system (Sartorius). (B) IncuCyte Images of iNs at 0, 4- and 8-days post plating with overlaid neurite (blue) and nucleus (magenta) segmentation masks (scale bar = 50 µm). (C) The results for cumulative neurite length. 16p11.2 deletion (16pDel) and duplication (16pDup) iNs showed significant differences in neurite length compared to WT and KCTD13 heterozygous deletion (KCTD13Het) at the p<0.05 level (one-way ANOVA). Shaded area indicates SEM. (D) Similar results for cumulative neurite branchpoints. 16pDel and 16pDup iNs showed significant difference in neurite branchpoints comparable to WT and KCTD13Het at the p<0.05 level (one-way ANOVA). Shaded area indicates SEM. (E) Post hoc comparisons of neurite length using the Tukey HSD test. 16pDel and 16pDup neurons displayed significantly reduced neurite length compared to WT neurons (*** p < 0.001), while KCTD13Het neurons displayed neurite length comparable to WT (p > 0.05). (F) Post hoc comparisons of neurite branchpoints using the Tukey HSD test. 16pDel and 16pDup neurons displayed significantly reduced neurite branchpoints compared to WT neurons (*** p < 0.001), while KCTD13Het neurons displayed a comparable level of neurite branchpoints compared to the WT group (p > 0.05). The number of images per group were WT n=170, 16pDel n=118, 16pDup n=105, and KCTD13Het n=87.

### 16p11.2 RGD neurons exhibit altered electrophysiological features

To assess the electrophysiological features of the 16p11.2 RGD neuronal cultures, we measured spontaneous neuronal firing using multi-electrode arrays (MEAs), a non-invasive platform for simultaneous recording of electric signals from multiple electrodes to study electrophysiology in vitro (Figure 5A). We differentiated iNs and then replated 60k Day 5 iNs onto MEA plates, then continued differentiation and recorded their activity directly for a period of time (Figure 5A). Representative temporal raster plots illustrating timestamps of spikes over 1 min of continuous recording and overlaid representative waveforms are shown in Figure 5B-E, respectively, for WT, 16pDel, 16pDup and KCTD13Het iNs. One-way ANOVA indicated significant differences in neuronal activity (normalized weighted mean firing rate, F=4.28, p=8e-3, df=3), functional connections between neurons (normalized synchrony, F=4.69, p=5e-3, df=3), and functional networks (normalized oscillation, F=5.21, p=3e-3, df=3) across WT, 16pDel, 16pDup, and KCTD13Het lines. We observed a significant effect of 16p CNVs on neuronal activity (WT: mean=1, SE=0.09; 16pDel: mean=0.65, SE=0.05, p=1.02e-3; 16pDup: mean=0.65, SE=0.06, p=3.23e-3) (see methods), while WT and KCTD13Het (mean=0.85, SE=0.12) were not significantly different (p=2.23e-1) (Figure 5F). The neurons with 16p11.2 CNVs and KCTD13Het all displayed significantly reduced synchrony (WT: mean=1, SE=0.22; 16pDel: mean=0.52, SE=0.08, p=7.97e-3; 16pDup: mean=0.51, SE=0.08, p=8.54e-3; KCTD13Het: mean=0.43, SE=0.05, p=1.13e-2) (Figure 5G). WT and KCTD13Het neurons were not significantly different (WT: mean=1, SE=0.05; KCTD13Het: mean=0.91, SE=0.06, p=2.76e-1) but 16pDel and 16pDup neurons exhibited significantly reduced oscillation (Del: mean=0.77, SE=0.04, p=7.39e-5; Dup: mean=0.78, SE=0.04, p=5.1e-5) (Figure 5H) when compared to WT. Like the morphometric phenotyping, these data suggest that changes in dosage of 16p11.2 genes affects the electrophysiological properties of iNs.

**Figure 5.**
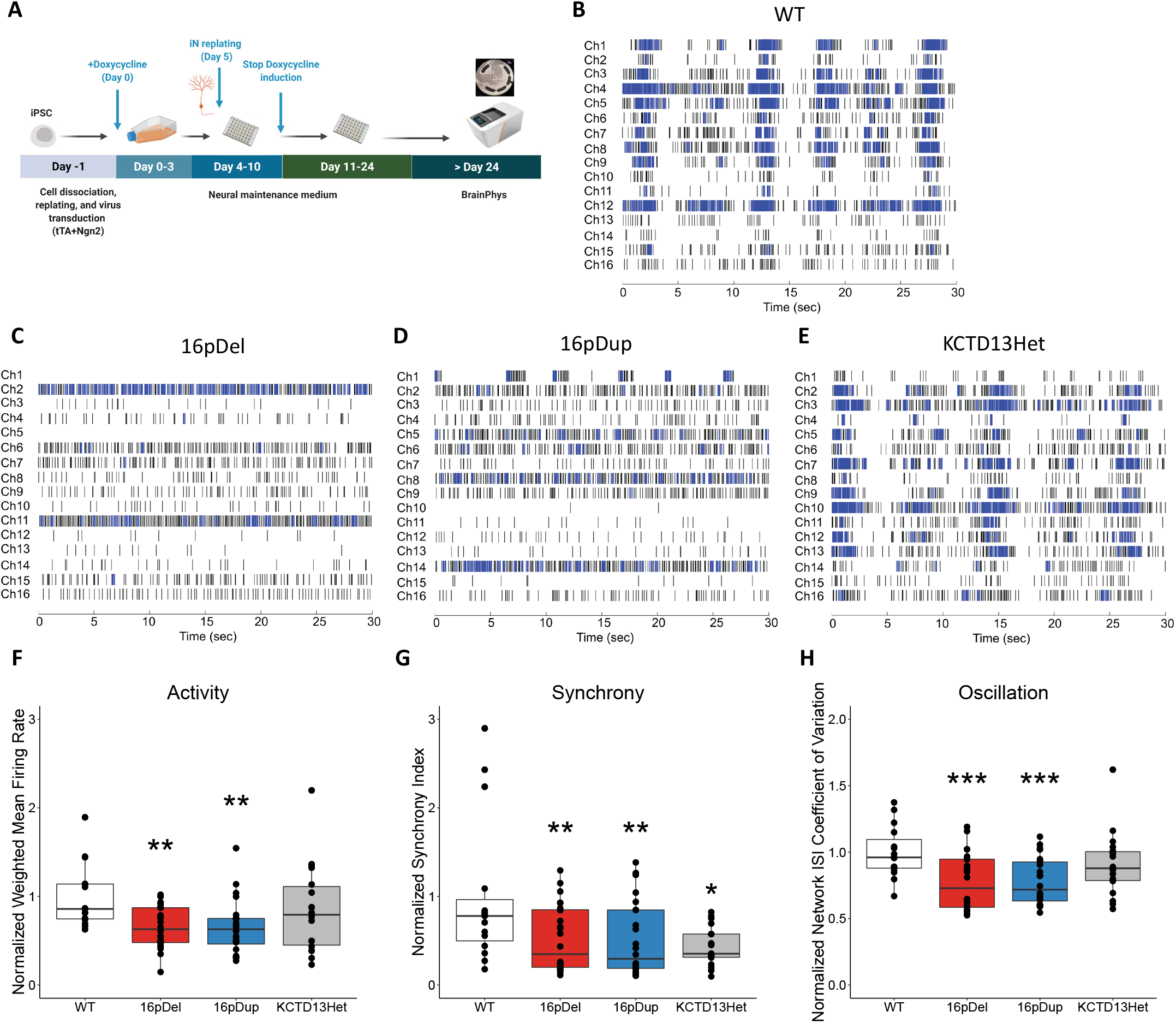
RGD neurons displayed aberrant electrophysiological properties. (A) Overview of study design. iNs were differentiated from hiPSCs as described in Materials and Methods. On Day 5 iNs were replated onto MEA plates with NMM. The neural activities were recorded after the culture medium switched to BrainPhys Neuronal Medium (Day 24). (B-E) Representative temporal raster plots from iN models demonstrating the activity over time for all electrodes in the well. Each plot is 30 s for sufficient spike and burst resolution, and horizontal rows correspond to 16 channel/electrodes; wild type (WT), 16p11.2 deletion (16pDel), 16p11.2 duplication (16pDup), KCTD13 heterozygous deletion (KCTD13Het). Raster plots generated with Neural Metric Tool v3.2.5 software (Axion Biosystems) (F) Neuron activity (normalized Weighted Mean Firing rate). 16pDel and 16pDup neurons displayed significantly lower activity compared to WT neurons (** p < 0.01), while KCTD13Het neurons displayed a level of activity comparable to WT (p = 0.222). Data are presented as mean ± SEM with normalized data points plotted. (G) Neuron synchrony (normalized Synchrony Index). 16pDel, 16pDup, and KCTD13Het neurons displayed significantly lower synchrony compared to WT neurons (* p < 0.05, ** p < 0.01). Data are presented as mean ± SEM with normalized data points plotted. (H) Neuron network oscillation (normalized Network ISI Coefficient of Variation). 16pDel and 16pDup neurons displayed significantly lower oscillation compared to WT neurons (*** p < 0.001), while KCTD13Het neurons displayed a level of activity comparable to WT (p = 0.276). The number of samples per group was WT n=15, 16pDel n=24, 16pDup n=24, and KCTD13Het n=18. Data are presented as mean ± SEM with normalized data points plotted.

### Altered cell complement in 16p RGD cerebral organoid model

The observation of an impact on the development and function of iNs by both 16p11.2 deletion and duplication, coupled with the fact that a number of the significant WGCNA modules from mouse brain and human NSCs were enriched for members of a module defined in early human neurodevelopment (‘M2’ in Figure 3), prompted us to assess potential neurodevelopmental deficits in cerebral organoids. We differentiated a subset of hiPSC lines with 16p CNVs or KCTD13 heterozygous inactivation into cerebral organoids using the protocols as described (Lancaster and Knoblich, 2014) and (Quadrato et al., 2017) and performed scRNAseq on 6-month-old (6M) organoids (n = 2 WT, 2 16pDel deletion, 2 16pDup duplication, and 2 KCTD13Het) to investigate genotype-specific single-cell signatures (Figure 6A). These data were analyzed by uniform manifold approximation and projection (UMAP), GO enrichment and co-expression analysis (WGCNA). Expression of canonical marker genes identified excitatory neurons, inhibitory neurons, and astroglia as three major cell classes in the cerebral organoids (Figure 6B and Figure S4A). A fourth cell population which did not pass criteria and was annotated as ‘unknown’ expressed limited oligodendrocyte and microglia-related markers (Figure 6B and Figure S4A). The cerebral organoids carrying 16p11.2 CNVs displayed reciprocally altered ratios of excitatory to inhibitory neurons in comparison to WT (Figure 6C), with 16pDel and 16pDup organoids having relatively more inhibitory and excitatory neurons, respectively. The KCTD13Het organoids showed no dramatic cell ratio changes compared to WT, again suggesting that the observed functional changes are a result of the combinatorial effects of many genes in the region and that the altered excitatory/inhibitory neuron ratio seen with the 16p11.2 deletion is not driven by KCTD13 alone.

**Figure 6.**
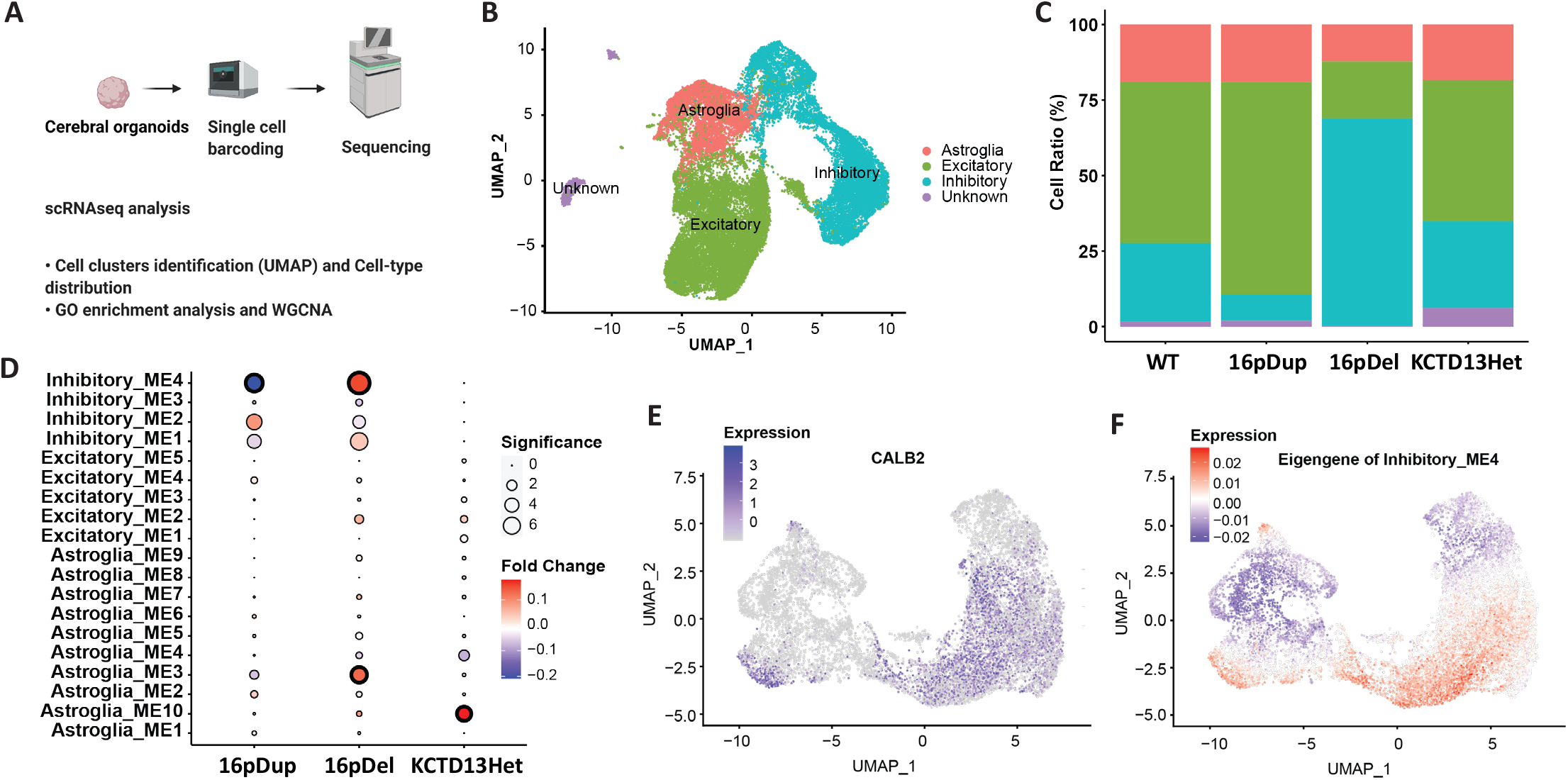
Altered neurodevelopmental signatures in 16p11.2 cerebral organoids. (A) Experimental design. 16p11.2 organoids were differentiated from hiPSCs as described in Materials and Methods. The organoids at 6 months were dissociated as single-cell suspensions and further processed by 10x Chromium and sequenced by illumina NovaSeq S4 platform. (B) Clustering of organoid cells in the UMAP space, with cell types assigned. (C) Proportion of cells in each cell type per genotype (WT, 16pDup, 16pDel and KCTD13Het). (D) Modules’ average expression change in each genotype compared to WT. The circle size represents the significance of expression changes in terms of negative-log10-transformed Bonferroni-corrected p values, while the color gradient represents the strength of expression changes in log-transformed scale. In inhibitory neurons, Inhibitory_ME2 contains all the six HVGs from the 16p11.2 region and is positively correlated with 16p11.2 CNV dosage as expected. (E-F) Normalized expression of *CALB2* (E) and the eigengene expression of Inhibitory_ME4 (F) across the inhibitory neuron population in the UMAP space.

To dissect the underlying molecular patterns that led to genetic lesions and cell-composition imbalance in these organoids, we investigated gene co-expression modules that are correlated with various 16p genotypes (Table S10). To avoid data sparsity from scRNA, only highly variable genes (HVGs) in each cell population were used. Although two modules from astroglia, “ME3 and “ME10”, showed overall upregulation due to 16p11.2 deletion and to heterozygous *KCTD13* deletion, respectively, indicating the potential for non-neuronal effects, the most significant correlation with 16p11.2 gene dosage was found in the “ME4” module of inhibitory neurons. Notably, Inhibitory ME4 was negatively correlated with 16p11.2 CNV dosage (Figure 6D and Table S10) and showed modest enrichment for NDD genes (Figure S4B), supporting the importance of an effect on inhibitory neurons. Indeed, GO enrichment analysis of its module member genes revealed significance for terms relevant to cell morphogenesis, neurogenesis, and neuron differentiation (Figure S4C and Table S11), while SynGO enrichment analysis revealed no significant term. Moreover, a number of the top-10 genes most correlated with its eigengene were highly relevant to GABAergic inhibitory neuron function (Figure S4D and Table S12). Among these, *GAD2* is a GABA synthesizing enzyme, and *DLX2* and *DLX5* are GABA interneuron progenitor transcription factors (Al-Jaberi et al., 2015). We further explored which subtype of inhibitory neurons was represented by Inhibitory ME4 using six subtype markers, including *CALB1, CALB2, NPY, PVALB, SST*, and *VIP*. Interestingly, we found that the expression pattern of CALB2 (encoding calretinin) across the inhibitory neuron population (Figure 6E) was most similar to the expression pattern of the Inhibitory ME4 eigengene (Figure 6F), indicating that the majority cell type represented in the ME4 module is the calretinin-positive GABAergic inhibitory neuron. Thus, both neuronal ratios and gene expression changes in the organoid analyses point most directly to GABAergic inhibitory neurons and the associated excitatory/inhibitory balance as a target of disruption that could contribute to neurodevelopmental and cognitive deficits in both 16p11.2 deletion and duplication.

## DISCUSSION

The 16p11.2 RGD is associated with a variety of prominent neurodevelopmental and other phenotypes, including features that are shared (e.g., NDD, ASD, seizure), mirrored (e.g., macrocephaly / microcephaly, obesity / low body weight), or distinct (e.g., predisposition to neuroblastoma among deletion subjects, schizophrenia among duplication subjects) (D’Angelo et al., 2016; Maillard et al., 2015; Weiss et al., 2008). Each of these features is likely to be driven by haploinsufficiency or triplosensitivity for one or more genes or combinations of genes within the 16p11.2 region. Indeed, it is conceivable that each of the shared or mirrored phenotypes results from the pathway disruptions in both deletion and duplication individuals that ensue from altered expression of the same critical CNV region gene or set of genes. However, while mutational analysis of persons with NDD in the absence of 16p11.2 dosage change have pointed to several different genes in the CNV region as potential contributors, none has been identified as unequivocally causal. Consequently, we sought to gain insight into this RGD through the shared (both reciprocal and non-reciprocal) and distinct transcriptional changes associated with deletion and duplication of the entire region.

Our overall findings in both the mouse and hiPSC model systems indicate that the expression of genes in the CNV segment directly reflects their gene dosage, but that the same dosage change results in significantly altered expression of largely different sets of genes outside the CNV segment in different tissues and cells. There was more sharing of these altered non-CNV genes than expected by chance, particularly across the brain regions, but those genes that were shared represented a small minority of the total DEGs and only a yet smaller subset of these displayed a reciprocal effect (i.e., expression altered in opposite directions by deletion vs. duplication). However, GO enrichment analyses of the significantly altered genes, and particularly co-expression of genes whose expression was correlated with CNV dosage, provided evidence for several commonly disrupted biological processes across the mouse brain regions and human neurons. These results were most prominent for alterations in energy metabolism, mRNA metabolism, translation and protein targeting. These alterations were not observed in the peripheral tissues, which exhibited disruption of distinct biological processes in each case.

Interestingly, although the vast majority of DEGs did not display a significant reciprocal effect of deletion and duplication on their expression, those WGCNA co-expression modules that correlated with CNV dosage were revealing of similar GO terms across the brain and human cell analyses, indicating that this enrichment is driven by more subtle reciprocal alterations of many genes involved in these biological processes. However, some significant co-expression modules, particularly the cerebellar module CE1, suggest the existence of additional interconnected effects on a large set of genes that shows limited GO enrichment, but which may be important in contributing to abnormal phenotypes. Expression of the genes in CE1 is positively correlated with CNV dosage, with an apparently larger effect of deletion than duplication, and boasts strongly significant enrichment for loss of function constrained genes and FMRP target genes, along with enrichment for NDD genes, DDD genes, chromatin modifiers and genes from a co-expression module defined very early in human neurodevelopment (M2). However, unlike the extensive sets of GO enrichments for most other significant mouse brain modules, CE1 is most significant for the term homophilic cell adhesion via plasma membrane adhesion molecules, with weaker support for other terms related to neuronal development (e.g., neuron differentiation; neuron development; neurogenesis) and adhesion (e.g., cell-cell adhesion via plasma membrane; biological adhesion) (Table S9). This contrast indicates that the large CE1 module and modules CO1, CO2, S3, and S5, which did not show any strong GO enrichments, might each reflect convergence of 16p11.2 dosage-elicited expression changes for subsets of non-CNV genes drawn from multiple different biological processes or could result from a diversity of 16p11.2 dosage-elicited responses in the different cell types within each region.

Together, the notable similarities and many differences in transcriptomic alterations between brain regions, peripheral tissues and human neuronal cells indicate that the CNV dosage changes of the same set of genes have impacts, both in terms of biological processes disrupted and in terms of the genes most significantly altered within those processes, that are context-specific. The importance of cellular context is reinforced by differences in the functional impact of the 16p11.2 CNV in our iNs (reduced numbers of neurites and branchpoints and electrical activity due to both deletion and duplication) compared to dopaminergic neurons (larger soma and hyperexcitability due to deletion and increased number of neurites due to duplication) derived from the same hiPSCs (Sundberg et al., 2021). Consequently, it is likely that multiple CNV genes and consequently disrupted biological processes contribute to the phenotypic features of the 16p11.2 RGD, potentially through a combination of different effects in different contributing cell types. For example, the cortex, cerebellum, and striatum are associated with a variety of functions, including movements, motor behaviors, learning, cognition functions and they are also implicated in various neurological diseases (Milardi et al., 2019). Recent data have shown that the cerebellum, known for sensory-motor control, also plays a role in social cognition and emotion due to its cortical connections. Consequently, the wide range of symptoms represented on the autism spectrum could result from the disruption of circuits involving all three of these brain regions (Crippa et al., 2016). Our findings suggest that the cortex-striatum-cerebellum network could suffer distinct disruption at each of its nodes, even if through different impacts on similar biological processes, that may all contribute to some degree to the ultimate neurodevelopmental phenotypes. Similarly, any phenotypes initiated in the periphery are extremely likely to be due to different biological processes than those critical to neurological phenotypes.

The biological processes that were most commonly disrupted due to 16p11.2 / 7qF3 CNV in the context of brain or cultured neuronal cells were energy metabolism, mRNA metabolism, translation and protein targeting. Although a number of the genes in the 16p11.2 CNV segment could participate in or impact on these processes, it is difficult to point to one as individually responsible for any of these disruptions. On the other hand, a number of the genes in the region have been implicated, primarily through model systems, as having potential impact on neurodevelopmental processes. These include, for example, *SEZ6L2* (synapse numbers, dendritic morphology, and neuritogenesis) (Qiu et al., 2021; Yaguchi et al., 2017), *CORO1A* (filopodia formation, required for initial neurite formation) (Alvarez Juliá et al., 2016; Dent et al., 2007), *DOC2A* (spontaneous neurotransmission associated with its calcium-dependent translocation) (Courtney et al., 2018; Groffen et al., 2006), *TLCD3B* (altered composition and levels of sphingolipids and glycerolipids associated with cellular membranes leading to synaptic protein mislocalization) (Tomasello et al., 2021), *TAOK2* (brain size, neural connectivity and excitatory transmission) (Richter et al., 2019); *PRRT2* (neuronal excitability) (Fruscione et al., 2018); and *MAPK3* (dendritic alterations of cortical pyramidal neurons) (Blizinsky et al., 2016). Recently, integration of genetic regulation of gene expression with genome-wide association data from human cohorts pointed to *INO80E* as the potential driver of schizophrenia due to 16p11.2 duplication and to both *SPN* and *INO80E* as contributors to the increased body mass index due to 16p11.2 deletion (Vysotskiy et al., 2021). Other model studies have also pointed to interaction between 16p11.2 genes as potentially being responsible for abnormal phenotypes (Iyer et al., 2018; McCammon et al., 2017). Consequently, our transcriptomic studies suggest that the various phenotypes observed in the 16p11.2 RGD likely result from both individual genes and gene interactions operating within the context of different cell types.

In the context of human glutamatergic iNs, the net functional effects of 16p11.2 deletion and duplication were similar in terms of neuronal morphometry and electrophysiology, yet in the context of cerebral organoids, these lesions produced reciprocal effects with respect to the content of GABAergic inhibitory neurons. Perhaps the importance of cellular context and specificity of phenotype is best exemplified by *KCTD13*. Based upon modeling in zebrafish (Golzio et al., 2012), this gene has been implicated previously as the source of the macrocephaly and microcephaly phenotypes associated with 16p11.2 deletion and duplication, respectively. Recently, RhoA and the associated Rho GTPase signaling pathway has been implicated in the link between *KCTD13* and 16p11.2 RGD phenotypes (Escamilla et al., 2017; Lin et al., 2015; Martin Lorenzo et al., 2021). However, in our human iN model system, *KCTD13* did not show haploinsufficiency for the neurite outgrowth and branching phenotypes and was minimally different from WT for the electrophysiological measures. Similarly, in cerebral organoids, *KCTD13* did not show a disrupted ratio of inhibitory vs. excitatory neurons. Consequently, *KCTD13* does not appear to be a contributor to these phenotypes in this context, but this does not preclude its contribution to the RGD phenotypes via other cellular or developmental contexts. Notably, in our co-expression analysis, cerebellum module CE1 and striatum module S1 yielded significant enrichment for the Rho GTPase cycle but module iN1 and the cortex modules CO1-4 did not (Escamilla et al., 2017; Lin et al., 2015; Martin Lorenzo et al., 2021), suggesting that the contribution of *KCTD13* dosage is limited to 16p11.2 RGD phenotypes involving cerebellum and striatum.

In the future, the organoid model approach may offer the best available experimental route to assess the effects of the 16p11.2 CNV and its constituent genes on human-relevant phenotypes that are generated through interaction of cell types in a developmental context. For example, in a 16p11.2 cohort study, the differences in intracranial volume, gray and white matter volume, cortical thickness and surface area in CNV carriers affected regions known to exhibit structural abnormalities in ASD and SCZ (Maillard et al., 2015). Moreover, immunohistochemical studies of both ASD and SCZ brains have showed a decrease in the caudate nucleus of the density of calretinin-positive interneurons (Adorjan et al., 2017, 2020). Genome-wide genetic analyses in humans have also supported the long-standing hypothesis that ASD involves disruption of the excitatory-inhibitory balance (Rubenstein and Merzenich, 2003; Satterstrom et al., 2020; Sohal and Rubenstein, 2019). Our data from cerebral organoids indicating dose-dependent differences in the ratio of excitatory to inhibitory neurons caused by the 16p11.2 CNV are consistent with this hypothesis and this system may offer a route to further dissect the pathways underlying this difference. By contrast, it has been reported that CALB2-positive GABAergic inhibitory neurons show greater abundance in adult primates than in rodents (Boldog et al., 2018; Wonders and Anderson, 2005). In addition, there are other subtypes that are numerous in primates, but are missing or greatly reduced in mouse cortex (Ballesteros Yáñez et al., 2005; Boldog et al., 2018). Therefore, fundamental differences in species context could potentially preclude the discovery of phenotype-associated signatures that are absent in the brains of model organisms.

In summary, we have demonstrated using isogenic hiPSC-derived neurons and mouse models that transcriptomic, morphological, electrophysiological and cell fate signatures of the 16p11.2 CNV are highly context dependent. While they provide evidence for disruption of a number of critical processes, most notably energy metabolism, mRNA metabolism, translation and protein targeting, and for the disruption of neuronal development and function, the details of these shared pathway disruptions vary by brain region, tissue and cell type. The shared pathway disruptions are also accompanied by alterations that are brain region, tissue or cell type-specific. Delineation of the individual combinations of causal or gene interaction and causal dosage change (deletion, duplication or both, either reciprocal or shared) presents a complex problem that will likely require a multi-faceted approach. However, our work suggests that the human cerebral organoid could be a particularly valuable tool for exploring human neurodevelopmental phenotypes of 16p11.2 RGD, as well as ASD and SCZ more generally (Battaglia et al., 2009; McCarthy et al., 2009).

## METHODS & SUPPLEMENTARY INFO

Detailed methods and supplementary information for this manuscript have been provided in a separate document, which will be linked directly from *bioRxiv*.

## ACKNOWLEDGMENTS

We thank S. Haggarty and S. Sheridan (Center for Genomic Medicine, Massachusetts General Hospital) for generously providing the control hiPSC line. These studies were supported by funding from the Simons Foundation for Autism Research (SFARI 328656 and 573206 (M.E.T.) and SFARI 308955 (J.F.G.)), the Nancy Lurie Marks Family Foundation (J.F.G. and M.E.T.), the US National Institutes of Health (NS093200, HD096326, MH115957, HD104224, MH123155, and GM061354 to J.F.G. and M.E.T.; MH123804 to RP), and Autism Speaks (J.F.G.). C.E.D.E. was recipient of a Rubicon Fellowship from the Netherlands Organization for Scientific Research (NWO).

## AUTHOR CONTRIBUTIONS

M.E.T., J.F.G., and D.J.C.T. conceived and designed the studies. D.J.C.T., J.W., K.M., and C.E.D.E. performed molecular studies. D.J.C.T. performed cerebral organoid differentiation. S.E., P.R., D.G., R.L.C., T.A., R.Y., and A.R. performed computational and statistical analyses. B.B.C., K.O., A.S., and N.D.B. performed RNAseq and single-cell RNAseq library preparations. E.M., W.M., D.L., C.E.D.E, A.R. and R.J.K. obtained mouse brain samples. A.H. performed optical genome mapping studies. D.J.C.T., C.E.D.E., J.W., R.P., M.E.T., and J.F.G. interpreted molecular datasets. D.J.C.T., S.E., D.G., P.R., X.N., M.E.T. and J.F.G. wrote the manuscript, which was revised by all authors.

