## Supplementary material for "Tissue and cell-type specific molecular and functional signatures of 16p11.2 reciprocal genomic disorder across mouse brain and human neuronal models": Methods and Supplementary figures

Derek J.C. Tai[1-4*], Parisa Razaz[2-4*], Serkan Erdin[1,2,4*], Dadi Gao[2-4*] et al.
*Full list of authors is available in the main text*

### Materials and Methods

#### CRISPR/Cas9 genome editing and cell model development

##### Guide RNA design, hiPSC culture, and DNA transfection

16p11.2 CRISPR-engineered, isogenic hiPSC lines with deletion or duplication of the 16p11.2 region were generated using the SCORE approach (Tai et al., 2016). Briefly, to design the optimal guide RNA, we first identified all possible 18–25mer guides with Jellyfish and performed a degenerate BLAST search to identify sequences that would uniquely target the 16p11.2 SDs, respectively, with no predicted off-target effects. The gRNA was cloned into pSpCas9(BB)-2A-Puro plasmid with a puromycin resistance marker (pX459, Addgene plasmid 48139) using a BbsI restriction site. Validation of the guide sequence in the gRNA vector was confirmed by Sanger Sequencing. Before transfection, all plasmids were purified using the EndoFree Plasmid Maxi Kit according to the manufacturer's instruction (Qiagen).

KCTD13 CRISPR-engineered, isogenic hiPSC lines with deletion of KCTD13 were generated via transfection with CRISPR guide RNA 5'-TAAAAAGGATGGATGTAGGC-3' and 5'-TGCCTGTGTTAGGAGGTATC-3' using the Amaxa Nucleofector II (Lonza) with Human Stem Cell Nucleofector Kit 1 (Lonza) and program B-016, according to the manufacturer's instructions. After nucleofection, hiPSCs were cultured in media supplemented with 10 μM Y-27632 dihydrochloride for 24 h prior to selection with puromycin (0.1 μg/mL). After 24 h, surviving hiPSCs were recovered in fresh Essential 8 medium for 48 h prior to FACS.

All hiPSCs were maintained in feeder-free culture on Matrigel hESC-Qualified Matrix-coated plates (Corning, 08-774-552) with Essential 8 media (Gibco, A1517001) supplemented with penicillin-streptomycin (Life Technologies, 15140) in a humidified incubator at 37°C with 5% $CO_2$. ReLeSR (STEMCELL Technologies, 05873) was used for routine cell passaging. mFreSR freezing medium (STEMCELL Technologies, 05855) was used for cryopreservation. Y-27632 dihydrochloride (MedChem Express, HY-10583) was added to media at 10 μM for up to 24 hours for initial plating and for several subsequent passages.

##### Single-cell isolation via fluorescence-activated cell sorting (FACS)

To obtain isogenic hiPSC colonies following CRISPR/Cas9 treatment, single cells were isolated via FACS. At 72 h after nucleofection, the hiPSCs were dissociated into a single-cell suspension using Accutase and resuspended in DPBS with 10 μM Y-27632 dihydrochloride (Santa Cruz Biotech). All samples were filtered through 5-mL polystyrene tubes with 35-μm mesh cell strainer caps (BD Falcon 352235) immediately prior to sorting. After adding TO-PRO-3 viability dye (Invitrogen), live (TO-PRO-3-) GFP+ hiPSCs were sorted on the BD FACSAria II with a 100-μm nozzle under sterile conditions and plated at one cell per well onto Matrigel-coated 96-well plates. Once multicellular colonies were clearly visible (2–3 d after sorting), they were collected into individual wells of Matrigel-coated 96-well plates by manual picking. Once individual hiPSC colonies were available (~14 d after sorting), the genomic DNA from those colonies was characterized by copy number assay.

#### Copy number analysis and cell lines characterization: quantitative real-time PCR (qRT-PCR), chromosomal microarray analysis and optical genome mapping

Array-based comparative genomic hybridization (aCGH) was performed on the CytoScan HD array (ThermoFisher) according to the protocol provided by the manufacturer. The assay tests for imbalances (gains or losses) in the genomic DNA sample. This array platform contains ~2.7 million probes throughout the human genome, including 1,953,246 copy number probes and 743,304 SNP probes. A genomic imbalance is noted when six or more oligonucleotides show a minimum average log ratio of 0.25 for one-copy gains and –0.50 for one-copy losses; oligonucleotide information is based on the human genome reference build NCBI 37.3 (hg19). It covers >36,000 RefSeq genes with 1 marker per 880 bases, complete ISCA constitutional coverage (1 marker / 384 bases), cancer gene coverage (1 marker / 553 bases), X chr genes (1 marker / 486 bases), and 12,000 OMIM genes (1 marker / 659 bases). Genomic imbalances are called using ChAS software (ThermoFisher) when a minimum of 50 consecutive probes is observed for loss, and 50 consecutive probes are observed for gain. A 50 Kb size cutoff (i.e., the lower limit of detection) has been established for CNV calls from constitutional specimens at the Jackson Laboratory. This assay does not exclude chromosome anomalies smaller than the assay's effective resolution. The assay is also not specifically designed to detect mosaicism, uniparental disomy, methylation abnormalities, or other chromosomal rearrangements (including chromosomal translocations, insertions, and inversions). For optical genome mapping, frozen hiPSC pellets were shipped to Bionano Genomics. Isolation of high molecular weight DNA, labeling, data assembly and identification of breakpoint regions were conducted by Bionano Genomics as described (Kriegova et al., 2021).

#### Selection and differentiation of TRA-1-60 Positive hiPSCs

Lines selected for differentiation underwent magnetic activated cell sorting (MACS) for expression of the TRA-1-60 cell surface marker for selection of pluripotent cells. Cells were separated using a MiniMACS Separator (Miltenyi Biotec, 130-090-312) with Anti-TRA-1-60 microbeads (Miltenyi Biotec, 130-100-832) following manufacturer instructions (~$2x10^6$ cells per line). TRA-1-60 positive cells were plated with Y-27632 dihydrochloride (10 μM), expanded, and cryopreserved using mFreSR. Cells within three passages of TRA-1-60 selection were used for differentiation into NSCs using the PSC Neural Induction Medium kit (ThermoFisher, A1647801) according to the manufacturer's protocol (MAN0008031). For all CRISPR lines, Passage 7 stage NSCs were dissociated for RNA-seq.

For differentiation to iNs, TRA-1-60 positive hiPSCs were plated as single cells at 80% confluence on a Matrigel-coated 6-well plate with Y-27632 dihydrochloride. Polybrene (hexadimethrine bromide; Sigma, 107689) was added at 8 mg/mL within three hours of re-plating. Cells were incubated with polybrene for 10-15 minutes prior to the addition of lentivirus. Lentiviral constructs for directed differentiation of hiPSCs into iNs were made as described previously (Zhang et al., 2013) and added to polybrene-treated hiPSCs. Cells were incubated with lentivirus for 24 hours, followed by a media change with regular E8. At least 48 hours following single-cell re-plating, transduced hiPSCs were cryopreserved and passaged for expansion. Transduced hiPSCs were expanded onto matrigel-coated T-25 flasks. Once all lines in a batch reached 70-80% confluence, cells were re-plated as single cells onto a new T-25 flask with Neural Maintenance Media (NMM) supplemented with Y-27632 dihydrochloride and 2 μg/mL Doxycycline (Clontech, NC0424034) to begin induction of TetO gene driving Ngn2 expression and Puromycin resistance (Day 0). The NMM we use in this study is adopted from Shi et al., 2012 (Shi et al., 2012). Twenty-four hours after re-plating, media was changed to NMM supplemented with 2 mg/L Doxycycline (Millipore Sigma, D9891) and 1 μg/mL Puromycin (Sigma), to begin selection of Ngn2-expressing cells (Day 1). Fresh

NMM with Doxycycline and Puromycin was added to cells to continue selection on Days 2 and 3. On Day 4, cells were detached using Accutase (ThermoFisher, A1110501) and re-plated onto Poly-L-Ornithine (10 µg/ml; Sigma-Aldrich, P4957) / Laminin (5 µg/ml; Sigma-Aldrich, L2020)-coated plates with NMM supplemented with 2 mg/L Doxycycline, 10 mg/L human BDNF (Pro-Spec, CYT-207), and 10 mg/L human NT-3 (PeproTech, 450-03). Cells were counted prior to re-plating using a Countess II Automated Cell Counter (Invitrogen, AMQAF1000) with $2.5 \times 10^5$ cells plated per well of a 12-well plate. Following re-plating, iNs were not exposed to air and required half-media changes every other day. On Day 6, fresh NMM with Doxycycline, human BDNF, human NT-3, and 2 g/L Cytosine β-D-arabinofuranoside (Sigma, C1768-100MG) to prevent glial growth. On Day 8, a half-media change with fresh NMM with Doxycycline, BDNF, and NT-3 was conducted. For subsequent media changes (Day 10+), NMM supplemented with only BDNF and NT-3 was added until cells reached Day 24 of differentiation, at which time cells were dissociated for RNA-seq.

#### Generation of cerebral organoids and dissociation for single-cell RNAseq

To generate cortical organoids from hiPSCs, we used TRA-1-60 positive hiPSCs with the protocol described (Lancaster and Knoblich, 2014; Quadrato et al., 2017). Briefly, embryoid bodies (EBs) were derived by dissociating hiPSC colonies and plating 9,000 single cells in each well of a 96-well ultra-low attachment plate (Corning 7007). On day 3, half of the media was replaced with embryoid body media without bFGF and Y-27632. On day 7, EBs were moved to 24-well low-attachment plates (Corning 3473). Media was changed every other day. On day 12, EBs were transferred to a droplet of Matrigel in 6-well low-attachment plates (Corning 3471). Media was changed every 3-4 days and organoids were moved to an orbital shaker placed in the incubator.

Single-cell suspensions of cerebral organoids were prepared using the Worthington Papain Dissociation System (Worthington Biochemical, LK003153) with previously described adjustments (Velasco et al., 2019). In short, cerebral organoids at 6 months were moved to 60 mm dishes, to which a solution of Papain/DNase was added. Using a new, sterile razor blade for each, organoids were minced to form < 1 mm pieces and incubated at 37°C for 30 minutes on an orbital shaker set to 70 rpm. Each sample was then mixed with a 1 mL pipette and returned for an additional 10-minute incubation. Cells were then triturated with a 10 mL pipette and transferred to new conical tubes to allow debris to settle. These suspensions were transferred to new tubes containing protease inhibitor solution, inverted to mix, and then passed through 40 µm cell strainers into new tubes which were centrifuged at 300 x g for 7 minutes. The resulting pellets were resuspended in DPBS with 0.04% BSA between 900-1,000 cells/µL and with viability ranging from 87-98%, according to the Countess II Automated Cell Counter. Additional wash/resuspension steps were omitted to promote cell viability.

#### Neurite dynamics measurement

Neurite dynamics measurement was performed using a live cell imaging system, Incucyte ZOOM system (Sartorius) with the automated IncuCyte NeuroTrack analysis platform. Time lapse images were acquired under incubated conditions at 37°C and 5% $CO_2$, as described (Song et al., 2018), but with some modifications regarding image analyses. Briefly, iNs were plated onto transparent 96-well plates at a density of 17,000 cells per well with IncuCyte NucLight Rapid Red Reagent (1:2000) and imaged every hour (9 image locations from each well) over 7 days at a resolution of 0.61 µm/pixel. The number of biologically independent lines per genotype in the experiment were WT n=3, 16pDel n=2, 16pDup n=3, and KCTD13Het n=2.

Phase contrast (cells) and red channel (nuclei) images (1392x1040 pixels) were segmented using Essen IncuCyte NeuroTrack software (2018A), and time course data for neurite length, neurite branchpoints, and number of nuclei were exported. The number of images analyzed per group were WT n=170, 16pDel n=118, 16pDup n=105, and KCTD13Het n=87. Further analysis was performed on these metrics using custom software written in Matlab (R2018b). Images with nuclear count less than 200 were omitted from analysis. Neurite length per nucleus and neurite branchpoints per nucleus were calculated by dividing neurite length and neurite branchpoints by the average number of nuclei detected during the first 8-12 h of the imaging period. These metrics were then smoothed using robust Lowess method with a window of 10 data points (20 h). Cumulative neurite outgrowth and branchpoints were computed using the sum of the first difference of the smoothed metrics. Outliers were removed by mutant group using median + 1.5*inter quartile range. Time course plots were generated using Gramm for Matlab (Morel, 2018). One-way ANOVA with Tukey post-hoc comparisons were used for statistical comparisons between mutant groups.

##### Microelectrode array (MEA) electrophysiology

To track spontaneous activity in neuronal cultures, 60k iNs (Day 5) were plated per well of a 48-well CytoView MEA plate (Axion BioSystems). NMM supplemented with only BDNF and NT-3 was added until cells reached Day 24 of differentiation as described above. On Day 24, iN culture medium switched to BrainPhys Neuronal Medium (STEMCELL Technologies) and then half of medium was changed every three days. Starting on Day 25, extracellular recordings were made on the Axion Maestro Pro machine (Axion BioSystems) to monitor spontaneous activity within the culture. All recordings were performed in BrainPhys and were started 5 minutes after the MEA plates were placed on the recording chamber for a duration of 15 minutes. The raw signals were acquired real-time and analyzed offline using Axion's Integrated Studio Navigator v3.4.1 software (Axion BioSystems). Measurements obtained by the manufacturer set thresholds include activity, electrode burst, network burst, synchrony, and oscillation metrics. To assess the strength of synaptic connections, Synchrony is measured as the Area Under the Normalized Cross-Correlation, a unitless measure between 0 and 1 (Halliday et al., 2006). A value of 1 means spikes are perfectly synchronous, whereas 0 indicates perfectly asynchronous. To assess the functional networks, Oscillation is a measure of how the spikes from all of the neurons in a well are organized in time as the coefficient of variation in inter-spike intervals for each electrode, averaged across electrodes in the well. High values indicate action potentials are not coordinated across neurons in the network. Spike raster plots were created in Neural Metric Tool v3.2.5 software (Axion BioSystems). The statistical results were created in Axion's Integrated Metric Plotting Tool v2.4.4 software (Axion BioSystems). MEA experiment was performed on two independent plates on different dates, named as two different batches. Data were analyzed from the wells with ≥8 active electrodes/well and normalized by corresponding wild-type (WT) mean within each batch. To test the statistical significance between groups (16pDel, 16pDup and KCTD13Het) and WT samples, multivariate linear model (~ edit + batch) was employed. The recordings from two 48-well MEA plates were included in the analysis based on the timepoint with highest average neuron activity shown in WT. The number of biologically independent lines for MEA analysis are WT n=2, 16pDel n=2, 16pDup n=2, and KCTD13Het n=2. The number of replicates per group were WT n=15, 16pDel n=24, 16pDup n=24, and KCTD13Het n=18 in the analysis reflected in Figure 5.

#### Transcriptomics

##### Mouse models for 16p RGD, samples, and underlying data sets

The 16p11.2 mouse models with reciprocal CNV of the syntenic 7qF3 region were created at the Cold Spring Harbor Laboratory by A. Mills and colleagues, as previously described in Horev et al., 2011 and provided by the Jackson Laboratories (stock numbers 013128 and 013129). Dissection of mouse tissues was performed simultaneously for all mice at 8 weeks of age. The mouse samples were split into five RNA extraction batches (labeled as B1-5) and three sequencing batches (labeled as DS1-3) (Table S1). An overview of the underlying data sets and samples used for these analyses is shown in Figure 1B, and more details can be found in Table S1. In brief, the following models and samples were analyzed. (1) 16p11.2 CNV mice. 350 RNAseq libraries from 101 mice with 16p11.2 CNV. In the initial 16 mice, we evaluated six tissues (liver, white fat, brown fat, cerebellum, striatum, and cortex), which enabled exploration of 16p11.2 tissue-specific effects; and in the 85 replication mice, we restricted our analysis to brain tissues (cortex, striatum, and cerebellum). (2) 16p11.2 CRISPR hiPSC-derived NSCs (n=28) and iNs (n=24). NSC samples were composed of two batches: Batch 1 (WT n = 6, 16pDup n=8) and batch 2 (WT n = 6, 16pDel n = 8).

#### Strand-specific RNAseq library preparation

All RNA samples were extracted with Trizol reagent according to the manufacturer's instruction (Invitrogen). RNA sample quality (based on RNA Integrity Number, RIN) and quantity were determined on an Agilent 2200 TapeStation and between 500-100 ng of total RNA was used to prepare libraries. 1 µL of diluted (1:100) External RNA Controls Consortium (ERCC) RNA Spike-In Mix (Thermo Fisher) was added to each sample alternating between mix 1 and mix 2 for each well in batch.

350 mouse RNASeq libraries were prepared with customized version of strand specific dUTP method (188 libraries) (Blumenthal et al., 2014; Levin et al., 2010) and TruSeq® Stranded mRNA Library Kit (Illumina) (162 libraries), while the same TrueSeq kit was used in the preparation of 52 human cell line RNAseq libraries (28 NSCs, 24 iNs). Both library preparation methods used polyA capture to enrich mRNA, followed by stranded reverse transcription and chemical shearing to make appropriate stranded cDNA inserts for the library. Libraries were finished by adding both sample-specific barcodes and adapters for Illumina sequencing followed by between 10-15 rounds of PCR amplification. Final concentration and size distribution of libraries were evaluated by 2200 TapeStation and/or qPCR, using Library Quantification Kit (KK4854, Kapa Biosystems), and multiplexed by pooling equimolar amounts of each library prior to sequencing. 350 RNASeq libraries were sequenced on multiple lanes of an Illumina HiSeq 2000-2500 platforms, generating median 38M paired-end reads of 50, 51 and 75 bp. 52 human cell line RNAseq libraries were sequenced on multiple lanes of an Illumina HiSeq 2500 platform, generating median 33.8M paired-end reads of 75 bp.

#### RNA sequencing quality control

Quality of sequence reads was assessed by fastQC (version 0.10.1) [Andrews, S. FastQC A Quality Control tool for High Throughput Sequence Data. bioinformatics.babraham.ac.uk projects/fastqc/, doi: citeulike-article-id:11583827]. Gene-based counts for mouse and human RNAseq libraries were generated by aligning sequence reads to the mouse reference genome, GRCm38 (v83) and the human reference genome, GRCh37 (v75) and relying on Ensembl gene annotations of these reference genomes using STAR (version 2.4.2a) (Dobin et al., 2013) with parameters "--outSAMunmapped Within --outFilterMultimapNmax 1 --outFilterMismatchNoverLmax 0.1 --alignIntronMin 21 --alignIntronMax 0 --alignEndsType Local --quantMode GeneCounts --twopassMode Basic". Quality of alignments was assessed by custom scripts utilizing Picard Tools (https://broadinstitute.github.io/picard/), RNASeQC (Graubert et al., 2021), and SamTools (Li et al., 2009). These quality checking assessments and exploratory analyses identified one outlier RNAseq in cerebellum samples, Dp835, one outlier in human NSC, H7 and five outliers in human iNs; Dels B5, H7, C3 and Dups B9, C5. These samples exhibited high duplication rate varying between 55-89% and low estimated library sizes between 3.4-20.5 M. Dp835 was also an outlier in other quality control metrics including exonic rate (0.41), intergenic rate (0.24) and intronic rate (0.35). Further exploratory analyses including clustering and principal component analyses (PCA) were implemented in R (version 3.4) using DESeq2 (version 1.18.1) (Love et al., 2014) and custom scripts. Exploratory analyses identified a striatum sample, t1992 as an outlier in batch 1 samples, which also exhibited high chimeric pair percentage (6.73%). These outliers were excluded from further analyses including differential expression and co-expression. Our analyses were restricted to mouse genes with 1-to-1 human orthologs, which were obtained from the Mouse Genomics Informatics database ( http://www.informatics.jax.org/homology.shtml) in July 2020.

#### CNV Region Expression Analyses

Genomics coordinates of engineered region in 16p11.2 deletion and duplication mouse models, spanning from *Slx1b* to *Sept1* were obtained from Horev et al., 2011. Since original coordinates were based on mm9 mouse reference genome, they were further converted to mm10 coordinates using Liftover UCSC (https://genome.ucsc.edu/cgi-bin/hgLiftOver), as chr7: 126,688,926-127,218,445. This region harbored 35 protein coding genes based on the Ensembl GRCm38 (v. 83) annotations: 27 human orthologous protein coding genes, two human segmental duplication genes *Slx1b* and *Bola2* , which were not duplicated in the mouse genome, and four additional genes (*Cd2bp2, Tbc1d10b, Mylpf, Sept1*), which were centromeric to the CNV and beyond human 16p11.2 CNV segment and a mouse specific gene *Gm42742* and *Pagr1b*, where the latter was removed in Figure 1A, since it was not a separate gene and has been removed in the most recent mouse genome. Breakpoints of human 16p11.2 CNV regions in our CRISPR/Cas9 treated cell lines were determined as Chr16: 29,487,574-30,226,919 based on human reference genome GRCh37 (v. 75), using CRISPR guides. This region harbored 32 protein coding genes based on the Ensembl GRCh37 (v. 75) gene annotation. In Figure 1A and D, we used the new gene name *TLCD3B/Tlcd3b* for a 16p11.2/7qF3 gene which was named as *FAM57B/Fam57b* in the reference genomes (GRCh37.v75, GRCm38.v83) used in our analysis. An uncharacterized gene, *RP11-37C12.3* (ENSG00000258130) was excluded from the region, since it was annotated as a pseudogene in the most recent human reference genome. To estimate the expression levels of SD genes *BOLA2B, BOLA2, SLX1A, SLX1B, SULT1A3* and *SULT1A4* in human cell lines, we generated two reference genomes in which genes at one SD region was masked using bedtools maskdata (Quinlan and Hall, 2010). This approach allowed better estimation of expression levels of unmasked SD genes. To this end, we first realigned the sequence reads to these reference genomes using STAR with above-described parameters. Next, we identified the sequence reads mapped to the SD genes' exons with no mismatch from which expression of a particular SD gene was estimated counting the fragments represented by two mated reads that were mapped to the same gene. Using these count data for six SD genes in the original raw count matrix, we converted raw count expressions of SD genes to transcript-per-million (TPM). Similarly, applying DESeq2 + SVA pipeline to the same count data matrix, we estimated fold changes of these six SD genes in deletion vs wild type and duplication vs wild type comparisons across human cell lines. Statistical significance of deviation of observed fold changes of SD genes from expected fold changes (0.75 for deletion; 1.25 for duplication) in each comparison was assessed by applying two-tailed one sample t-test.

#### Differential gene expression analyses

Differential expression (DE) analyses performed on genes that passed the expression threshold in a given comparison using R/Bioconductor packages DESeq2 (v.1.18.1) (Love et al., 2014). To determine genes that passed the expression threshold for a particular comparison, we first calculated count-per-million (cpm) expression values of genes across the samples used in the comparison. Cpm expression of $i^{th}$ gene in sample j was defined as 1e6 x $C_i$ / $LS_j$, where $C_i$ is raw counts of $i^{th}$ gene and $LS_j$ is the library size of $j^{th}$ sample. We used the total number of uniquely mapped reads reported by STAR for a given sample for the library size of that sample. Next, we calculated cpm expression threshold corresponding to 10 counts for the particular comparison using the equation 1e6 x 10 / median(LS), where median(LS) is the median value of library sizes of samples used in the comparison. Cpm thresholds varied between 0.27 and 0.38 across the comparisons for which we performed DE analysis. All the genes, regardless of their type (e.g., protein coding, antisense) with expression values in cpm equal or greater than the cpm expression threshold in at least 50% of samples in either condition (e.g., Del vs WT) were further analyzed in the DE analysis. We performed DE analysis for each mouse tissue/human cell line type. In these analyses, CNV type (deletion or duplication) was compared with CNV type matched wild type samples in mouse non-brain tissues (e.g., Del vs Del WT). When we compared deletion or duplication to their CNV matched wild types in brain tissues, we also added wild types not specific for a certain CNV type from the third batch (B3) or the second data set (DS2) to each of their corresponding wild types (e.g., Del vs WT + Del WT). Deletion and duplication samples were compared to the same wild types in human iNs, whereas deletion and duplication samples were compared to their separate WTs in human NSCs. To account for unknown sources of variation in the expression data, surrogate variables (SVs) were estimated for each comparison using R/Bioconductor package the Surrogate Variable Analysis (SVA version 3.26) by setting ~ genotype as the full model and ~ 1 as the reduced model (Leek, 2014; Leek et al., 2012). Differentially expressed genes (DEGs) were identified at False Discovery Rate (FDR) < 0.1 using the Wald test under the design: ~ genotype + SVs, where FDR was calculated following the Benjamini Hochberg procedure (Benjamini and Hochberg, 1995). In these analyses, DESeq2's independent filtering and cooksCutoff options were turned off. Protein coding DEGs were used in the further downstream analyses including enrichment analyses and comparisons of DEGs across different comparisons. Statistical significance of overlap of DEGs between different comparisons was assessed by employing one-sided Fisher's exact test or equivalently, Hypergeometric test.

#### Gene co-expression network analyses

Gene co-expression network analysis for each mouse brain tissue and human cell types was performed separately using R package Weighted Correlation Network Analysis (WGCNA version 1.61) (Langfelder and Horvath, 2008). For this analysis, we used log2 transformed SVA corrected counts under signed network option and setting minimum module size to 50 and merging modules with > 75% similarity. SVA corrected counts for each brain tissue and human cell lines were generated by combining deletion, duplication and wild-type samples after removing outlier samples that were described above. SVA was applied to the union of analyzed genes in deletion vs wild-type and duplication vs wild-type comparisons relying on the full model ~ genotype (deletion, duplication, wild-type) and the reduced model ~ 1. Genes in the 7qF3 engineered region and 16p11.2 region were excluded in the further co-expression analyses of mouse and human samples respectively. Negative SVA corrected counts were set to zero and log2 transformed after adding 1 to the count matrix. To identify additional outlier samples in co-expression analysis, we adapted the procedure described (Oldham et al., 2008). as follows. We first computed the average Pearson's correlation coefficient for each sample by correlating that sample with other samples within each mouse brain tissue and human cell type. Further, from these sample specific average correlation values, tissue or cell type specific average correlation values and standard deviations were computed. Within each mouse brain tissue or human cell type, samples with average correlation values at least three standard deviations lower than tissue or cell type specific average correlation values were identified as outlier samples. In doing so, we identified the following samples as outliers in co-expression analyses: D809 for mouse cerebellum, w1514 and w1516 for mouse cortex, w1210 for mouse striatum and C5 for human NSCs. These outlier samples were excluded from further analyses. Soft power was selected such that scale-free topology fit ($R^2$) > 0.85. Module membership for each was re-evaluated based on the module membership p_value such that genes with p ≥ 0.01 were marked as unassigned (grey module). To identify the modules highly correlated with deletion and duplication in each co-expression analysis, we investigated how expression patterns of module genes represented by module eigengenes correlated with four situations in which deletion, duplication and wild type samples were represented by the following contrasts: 1) dosage effect (-1, log2(3/2), 0), 2) genotype effect ("cnv","cnv","wt"), 3) deletion vs duplication + wt ("cnv","wt","wt") and 4) duplication vs deletion + wt ("wt","cnv","wt"). Statistical significance of relationship between module eigengenes and the above contrasts was assessed using linear regression models, $eigengene_i \sim vector_j$, where i and j index module in a particular mouse brain tissue or human cell type and one of four situations described above respectively. Modules with p < 1e-5 and size ≥ 30 protein coding genes were selected for further analyses. To profile the expression pattern of module genes across the human neurodevelopment, we generated an expression matrix for selected modules using the

PsychENCODE expression data (Li et al., 2018), and calculated module eigengenes for each module by employing WGCNA. Next, we calculated the mean module eigengene value for each of nine developmental windows per module as described (Li et al., 2018). Statistical significance of module overlaps was assessed by employing one-sided Fisher's exact test or equivalently, Hypergeometric test.

#### Overrepresentation analysis

To assign biological significance to selected protein coding genes including DEGs, DEGs shared between two or more comparisons and module genes identified in co-expression analysis, overrepresentation analysis (ORA) of these genes for Gene Ontology (GO) Biological Process terms (Ashburner et al., 2000), which were retrieved from MSigDB (v7.4) database (Subramanian et al., 2005), synaptic gene ontologies (SynGO version 1.1) (Koopmans et al., 2019), and curated gene sets was performed using one-tailed Fisher's exact test equivalent to hypergeometric test. ORA was performed only for protein coding genes, where both selected and background gene sets contained only protein coding genes. To calculate enrichment p-value for a particular selected gene set, analyzed protein coding genes from which selected genes were chosen constituted the background gene set. For example, if ORA would be performed for DEGs from deletion vs wild type comparison in mouse cortex tissue, then the background set was all the protein coding genes that were analyzed in this analysis, or in other words, all the protein coding genes that passed the expression threshold. Similarly, if ORA would be performed for a particular module identified in co-expression analysis of mouse cortex samples, then the background gene set was all the protein coding genes analyzed in this co-expression analysis. In case of ORA of selected genes shared between two or more comparisons, the background gene set was an intersection of background gene sets for each comparison. In these analyses, both human and mouse genes were represented by their symbol identifiers and 1-to-1 human orthologs were used for the latter group. Furthermore, only GO terms with at least 10 associated genes in the background set were considered.

#### Generation of single-cell RNAseq libraries

Dissociated cells were maintained on ice no longer than 30 minutes prior to loading onto the 10X Chromium Single Cell Controller (10x Genomics, PN-120263). scRNA-seq libraries were prepared using the Chromium Single Cell 3' Library & Gel Bead Kit *v3* (PN-1000075) according to manufacturer's instructions (CG000183) using a Chromium Single Cell B Chip (PN-1000073) for a targeted cell recovery of 5,000 cells. Post-library construction quality control and quantification were performed using both a High Sensitivity D1000 ScreenTape (Agilent, 5067-5582) and qPCR via the Universal KAPA Library Quantification Kit (Roche, KK4824). Final libraries were pooled according to molar concentrations and submitted for sequencing on the illumina NovaSeq S4 platform averaging 8 libraries per lane at 2.5 billion reads per lane.

#### Single-cell RNAseq analysis

Each scRNA library was aligned against Ensembl human transcriptome reference GRCh38 version 92 via 10X CellRanger (v3.0.2) program with 5,000 expected cells. Each aligned library was then processed in Seurat v4.0.0.9010 (Satija et al., 2015) independently by filtering out cells with highest and lowest 2.5% quantile RNAs. All filtered libraries were merged, followed by an unsupervised clustering via UMAP method (Becht et al., 2019). The number of cell clusters were determined with resolution of 0.2. The unsupervised cluster labels were transferred to annotated neuron cell types heuristically, based on the expression levels of the selected cell type markers. Empirically, Clusters 0, 4, 5 and 6 were labeled as excitatory neurons while Clusters 1 and 3 were labeled as inhibitory neurons. Cluster 2 was labeled as astroglia cells.

Excitatory neurons, inhibitory neurons and astroglia cells were then analyzed independently. Each of these three cell types was further unsupervised clustered via UMAP method to reveal its sub-cell types. The number of sub-cell clusters were determined with resolution of 0.2. Meanwhile, WGCNA v1.70.3 (Zhang and Horvath, 2005) was applied on highly variable genes from excitatory neurons, inhibitory neurons and astroglia cells, respectively, with soft power of 8, 7 and 5. In each cell type, raw modules were merged with a dissimilarity threshold of 0.25. To identify the genotype effect on each module, the average expression of genes associated with a module in a given genotype was compared to that in WT. Wilcoxon Rank Sum and Signed Rank test was implemented to determine the significance. Bonferroni correction was applied to p values from all identified modules in all the three cell types. To reveal whether a module was enriched for a certain group of genes, Fisher's exact test was implemented.

**Data and Software Availability**

The RNA-seq data generated for the chromosome 16p11.2 study have been uploaded to the National Database for Autism Research (NDAR) repository under collection number 2304.

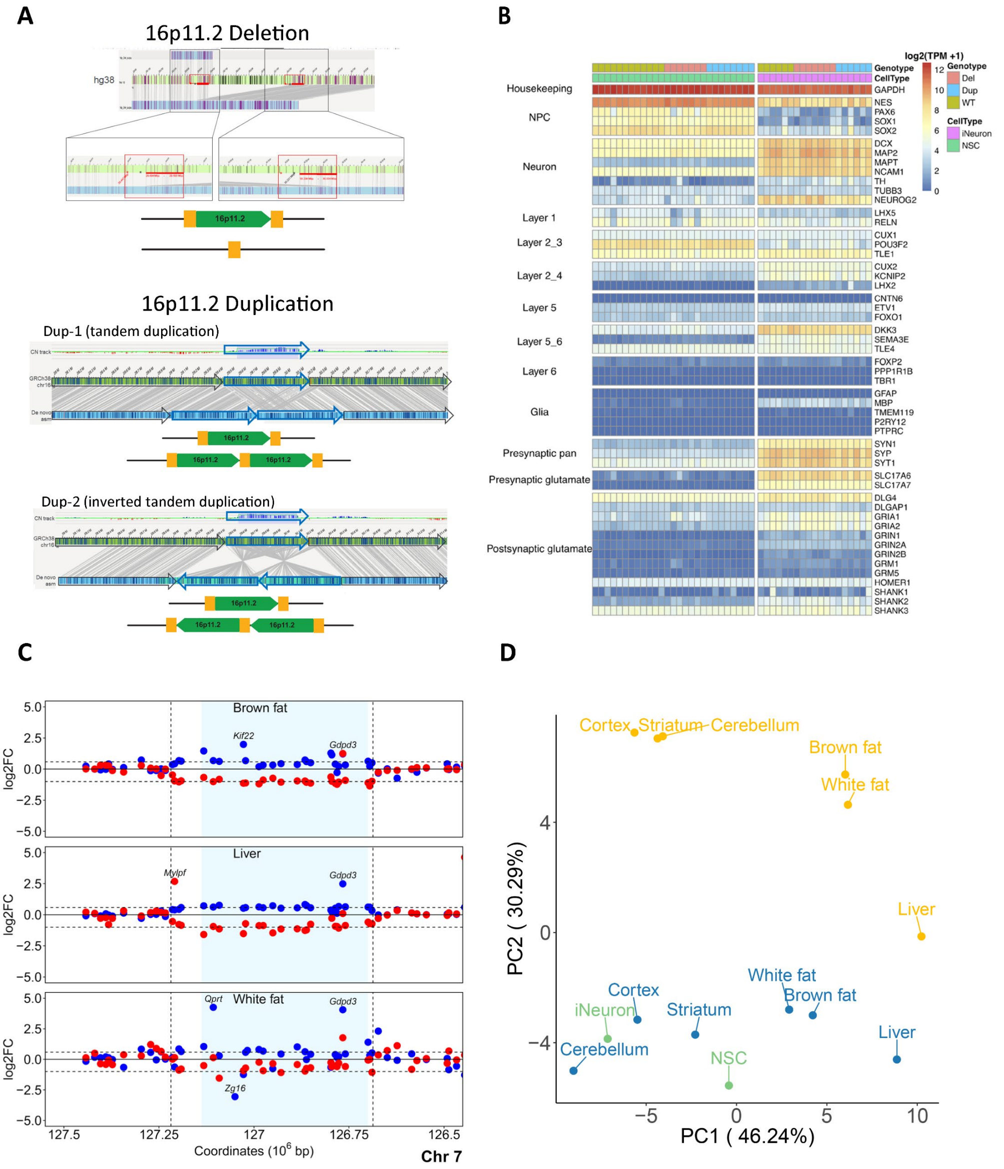

*Supplementary Figure 1:Characterization of hiPSC lines, neuronal derivatives, and mouse tissues*

(A) Chromosome architecture in hiPSC lines with 16p11.2 RGD. To further characterize genome architecture, nanopore sequencing and optical genome mapping was performed for a subset of 16RGD hiPSC lines (one deletion and the two duplication lines). Using optical genome mapping we can resolve chromosome architecture at 16p11.2 region. In the deletion line, the unique region has been deleted and only one copy of SD region left. In the duplication lines, the unique region and SD have been duplicated. Dup-1 is a canonical tandem duplication, and Dup-2 has an inversion of the tandem duplication. (B) Heatmap for cell type specific marker gene expression across all NSC and iN lines. (C) Fold change (log2) of the protein coding genes in the CNV and in the flanking regions are shown in coordinate space for deletions in red and duplications in blue across non-brain tissues. Light blue shaded region in the left panel highlights the unique portion of 16p11.2 CNV region harboring 27 human orthologous protein coding genes in the mouse 7qF3 segment. (D) PCA plot for all the samples based on the expression profile of the 27 CNV genes.

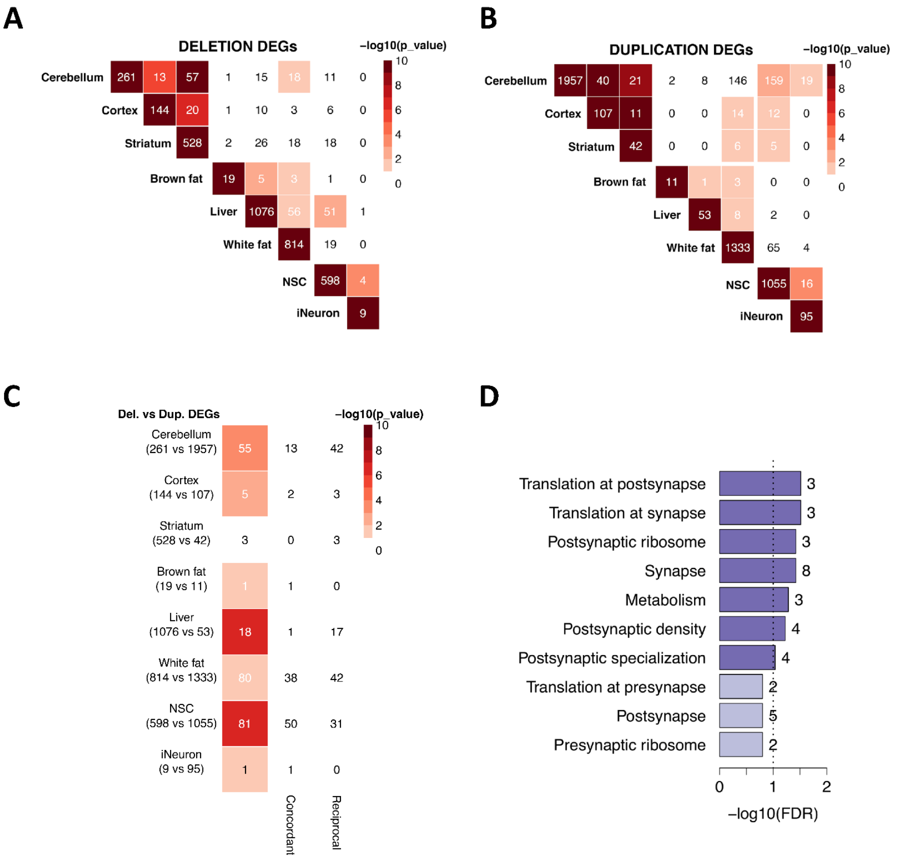

*Supplementary Figure 2: Shared DEGs and GO analysis*

(A) Overlap among the deletion DEGs observed between mouse brain tissues, peripheral tissues, and human cells. (B) Overlap among the duplication DEGs observed between mouse brain tissues, peripheral tissues, and human cells. (C) Significant sharing of Del and Dup DEGs among samples. Some shared DEGs have reciprocal responses to 16p11.2 CNV, and some shared DEGs consistently dysregulated in the same direction (concordant). (D) SynGO enrichment analysis for shared human NSCs and iNs DEGs.

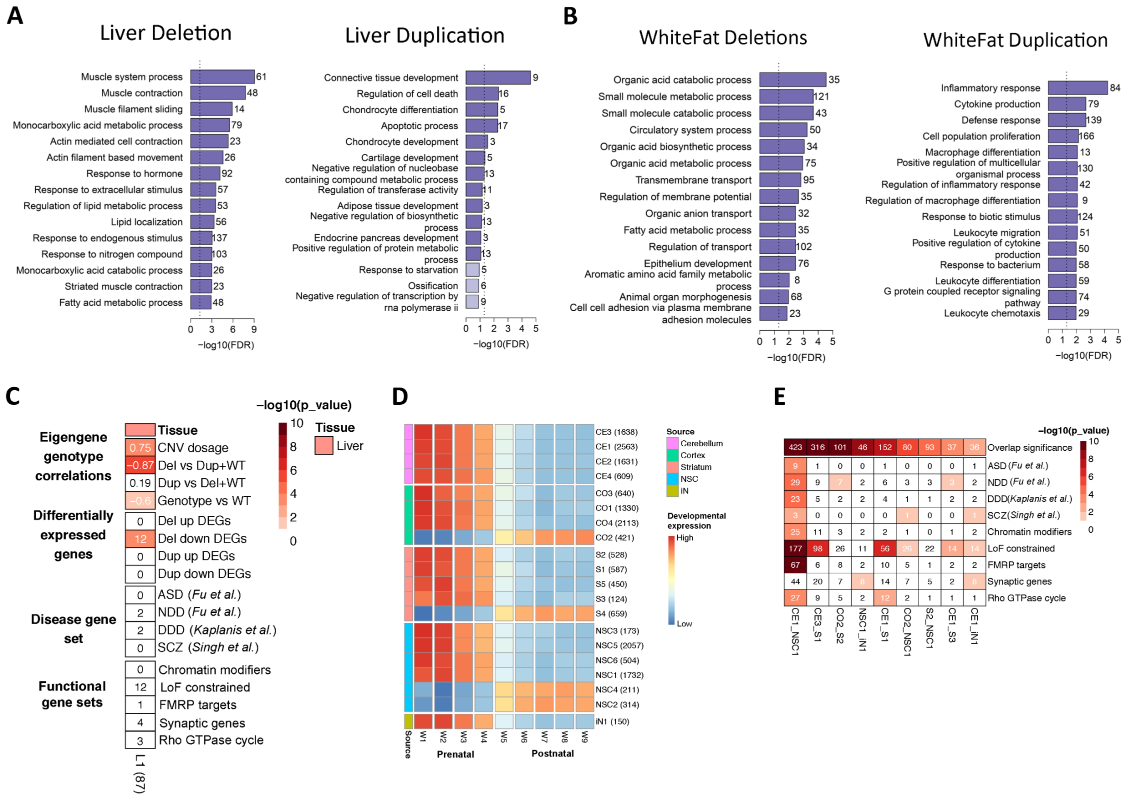

*Supplementary Figure 3: GO enrichment analysis for mouse non-brain tissues*

(A) *Left:* GO enrichment analysis for mouse liver deletion DEGs; *Right:* GO enrichment analysis for mouse liver duplication DEGs. (B) *Left:* GO enrichment analysis for mouse whitefat deletion DEGs; *Right:* GO enrichment analysis for mouse white fat duplication DEGs. (C) Modules that are statistically significantly associated with Del and Dup genotypes at p < 1e-5. The top panel shows eigengene genotype correlations, where numbers within the heatmap cells are Pearson's correlation coefficient. The second panel shows the statistical significance of overlap between up- and down-regulated differentially expressed genes (FDR < 0.1) from Liver deletion and duplication samples and liver module L1. The next two panels show module enrichment analyses against literature-curated gene lists; disease gene sets and functional gene sets. The lists include ASD associated genes(Fu et al., 2021), NDD associated genes(Fu et al., 2021), DDD associated genes (Kaplanis et al., 2020), rare variants in genes associated with schizophrenia (Singh et al., 2022), chromatin modifiers (Iossifov et al., 2014), loss-of-function intolerance constrained genes (LOEUF < 0.35) as reported by the genome aggregation database consortium (Karczewski et al., 2020), FMRP targets (Darnell et al., 2011), and synaptic genes from SynGO v1.1 (Koopmans et al., 2019). Numbers within the heatmap cells are number of genes shared between selected gene set and liver module, L1. Numbers within the parentheses show the number of protein coding genes in the co-expression modules identified in the mouse liver tissue. (D) The expression pattern of co-expression modules across brain developmental stages. Developmental expression values are mean eigengene values calculated using the PsychENCODE data for a given window (W1-9). (E) The overlap of co-expression module genes and their enrichments for selected disease and functional gene sets. The first row in the heatmap shows the statistical significance of overlap of module genes in -log10 scale, while the other rows show the enrichment of module genes against gene sets in -log10 scale. Numbers in the cells are the number of genes shared between selected modules (the first row) and the number of genes shared between gene sets and module (other rows).

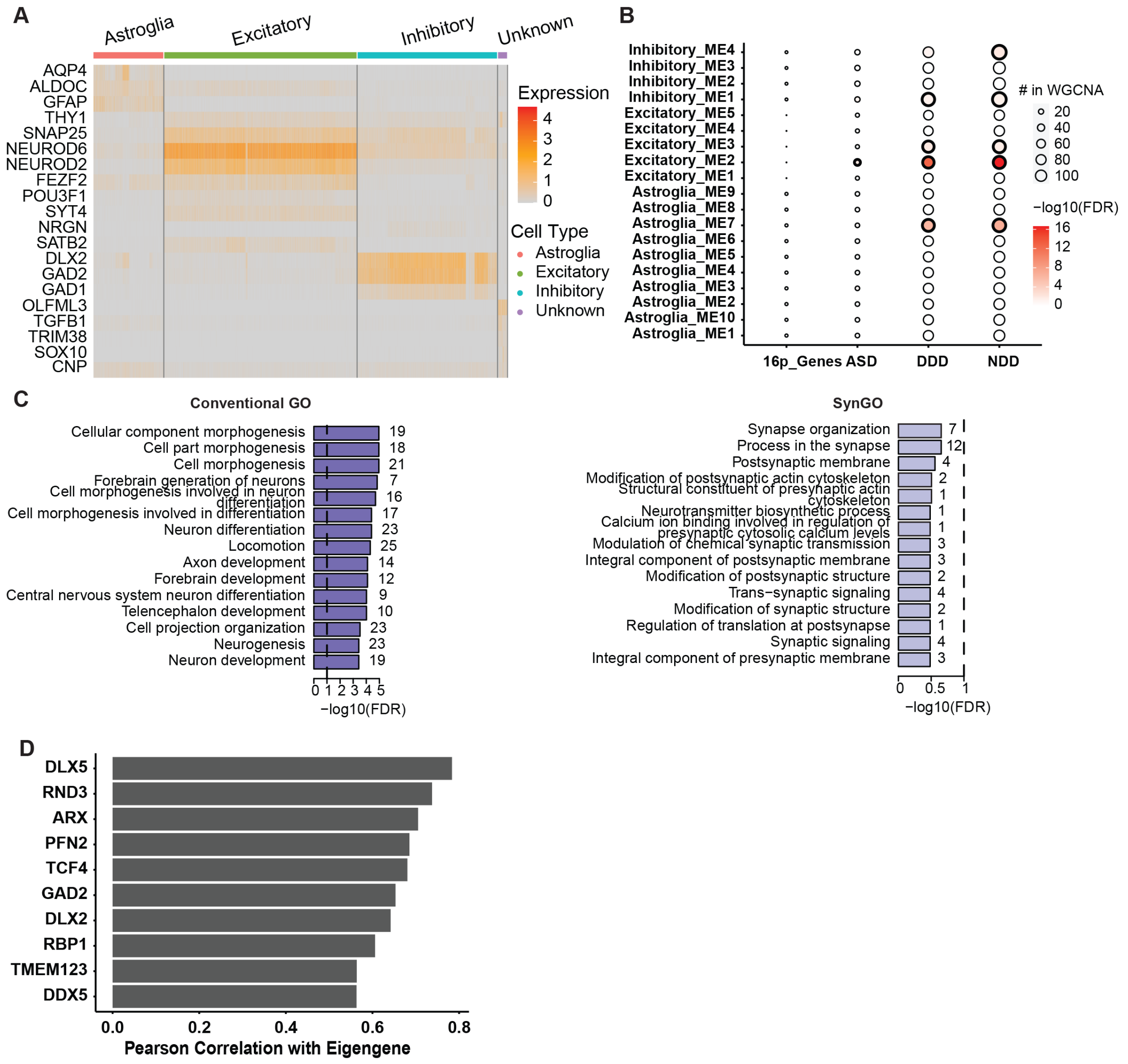

*Supplementary Figure 4: Transcriptomic signatures among models*

(A) Heatmap for canonical marker expression across organoid cells measured by scRNA. (B) Enrichment of each gene set in each of the identified cell-type-specific co-expression modules. The circle size represents the number of genes used in the WGCNA analyses (i.e., considered as highly variable genes), while the color gradient represents the significance of enrichment in terms of negative-log10-transformed FDR. (C) Conventional GO (*left*) and SynGO (*right*) enrichment analysis of genes in Inhibitory ME4. The vertical dashed lines indicate the cutoff for significance at FDR=0.1 in negative-log10-transformed scale. (D) Top 10 Pearson correlation coefficients of the correlation between the eigengene of Inhibitory ME4 and its gene members.

*Supplementary Table 1: Mouse data sets*

Distribution of mouse samples into three sequencing batches (datasets 1,2,3 (DS1-3)), five RNA extraction batches (B1-5), three genotypes (7qF3 deletion and duplication and wild types) across six mouse tissues: cortex (ctx), striatum (str), cerebellum (cbm), white fat (wfat), brown fat (bfat). Strand specific paired-end RNAseq libraries were prepared using a custom protocol adapted manually from Levin et al., 2010(Levin et al., 2010) and Illumina TruSeq. Red, blue and black colors highlight deletion, duplication and wild type samples respectively. Del-wt and dup-wt indicate wildtype samples extracted from control littermates matched with deletion and duplication liters respectively.

*Supplementary Table 2: Differential expression results of all genes/tissues/cells shown in Figure 1E and S1C*

Differential expression results as expression values in TPM, log2FC, p-values and FDR from genes in the CNV and flanking regions as shown in Figure 1E and S1C. (A) mouse tissues, (B) human NSCs and iNs.

*Supplementary Table 3: DEG counts from 16p RGD mouse and human NSC and iN models*

Number of differentially expressed protein coding genes from deletion vs wild-type and duplication vs wild-type comparisons across mouse brain and non-brain tissues as well as human cell lines at nominal $p < 0.05$ and FDR < 0.1. Differentially expressed genes (DEGs) were categorized into two respective groups: genes inside and outside the 16p11.2 (human)/7qF3 (mouse) deletion/duplication segment.

*Supplementary Table 4: Shared DEGs and the full list of GO terms*

(A) 223 unique DEGs shared by at least two brain regions, and the 28 unique DEGs shared between the NSC and iNs. (B) Gene ontology (Biological Process) terms enriched at nominal $p < 0.05$ for DEGs from deletion vs wild-type and duplication vs wild-type as well as their union (combined) across six mouse tissues and two human NSCs and iNeurons. P values of enriched terms for each comparison were adjusted applying Benjamini-Hochberg and Bonferroni procedures.

*Supplementary Table 5: The full list of SynGO terms*

SynGO terms enriched at nominal $p < 0.05$ for DEGs from deletion vs wild-type and duplication vs wild-type as well as their union (combined) across six mouse tissues and two human NSCs and iNeurons. P values of enriched terms for each comparison were adjusted applying Benjamini-Hochberg and Bonferroni procedures.

*Supplementary Table 6: The full list of differential expression analysis results and co-expression modules*

Differential expression analysis results of 16p11.2 (syntenic 7qF3) deletion vs wild-type and duplication vs wild-type comparisons and co-expression modules. (A) mouse tissues, (B) human NSCs and iNs.

*Supplementary Table 7: Table of module eigengenes*

Module eigengenes of all the modules identified by WGCNA for six mouse tissues; cerebellum (A), cortex (B), striatum (C), brown fat (D), liver (E), white fat (F), and human NSCs (G) and iNs (H).

*Supplementary Table 8: Correlation statistics of co-expression modules with various 16p genotypes*

Correlation statistics of co-expression modules identified in mouse tissues (A) and human NSCs and iNs (B) with various 16p genotypes. Modules were sorted by minimum p-values from four tests in ascending order within a tissue/cell type.

*Supplementary Table 9: Full list of GO and SynGO terms enriched for selected co-expression modules*

(A) GO Biological Process and (B) SynGO enrichment analysis results for selected modules from co-expression analyses of mouse tissues and human cell lines. Terms enriched at $p < 0.05$ are listed. Multiple testing correction was performed within each module (module_FDR, module_bonferroni) and across all the modules excluding grey module identified in a particular tissue/cell type applying Benjamini-Hochberg and Bonferroni procedures.

*Supplementary Table 10: Single-cell gene co-expression modules that are correlated with various 16p genotypes in cerebral organoids*

Two-sample Wilcoxon rank sum and signed rank test statistics for between average gene expression of WT and 16p genotypes for each cell-population-specific co-expression modules.

*Supplementary Table 11: Full list of GO and SynGO terms enriched for inhibitory ME4 module from scRNA co-expression analysis*

GO Biological Process and SynGO enrichment analysis results for ME4 module from scRNAseq co-expression analysis. Multiple testing correction was performed separately for GO and SynGO applying Benjamini-Hochberg and Bonferroni procedures.

*Supplementary Table 12: List of gene symbols in scRNA co-expression modules*

Gene symbols of genes in each of the cell-population-specific co-expression modules.
